# Altered plasma membrane abundance of the sulfatide-binding protein NF155 links glycosphingolipid imbalances to demyelination

**DOI:** 10.1101/2022.09.15.508082

**Authors:** Shannon J. McKie, Alex S. Nicholson, Emily Smith, Stuart Fawke, Eve Caroe, James C. Williamson, Benjamin G. Butt, Denisa Kolářová, Ondřej Peterka, Michal Holčapek, Paul J. Lehner, Stephen C. Graham, Janet E. Deane

## Abstract

Myelin is a multi-layered membrane that tightly wraps neuronal axons enabling efficient, high-speed signal propagation. The axon and myelin sheath form tight contacts, mediated by specific plasma membrane proteins and lipids, and disruption of these contacts causes devastating demyelinating diseases. Using two cell-based models of demyelinating sphingolipidoses, we demonstrate that altered lipid metabolism changes the abundance of specific plasma membrane proteins. These altered membrane proteins have known roles in cell adhesion and signalling, with several implicated in neurological diseases. The cell surface abundance of the adhesion molecule Neurofascin, a protein critical for the maintenance of myelin-axon contacts, changes following disruption to sphingolipid metabolism. This provides a direct molecular link between altered lipid abundance and myelin stability. We show that the Neurofascin isoform NF155, but not NF186, interacts directly and specifically with the sphingolipid sulfatide via multiple binding sites and that this interaction requires the full-length extracellular domain of NF155. We demonstrate that NF155 adopts an S-shaped conformation and preferrentially binds sulfatide-containing membranes in *cis*, with important implications for protein arrangement in the tight axon-myelin space. Our work links glycosphingolipid imbalances to disturbance of membrane protein abundance and demonstrates how this may be driven by direct protein-lipid interactions, providing a mechanistic framework to understand the pathogenesis of galactosphingolipidoses.

## INTRODUCTION

Myelination of axons enables fast, saltatory impulse propagation, facilitating increased neural processing speed and energetic efficiency (1). Myelin is a multi-layered membrane that is produced by oligodendrocytes in the central nervous system. Oligodendrocytes form numerous cellular extensions that wrap around axons to form distinct myelin internode segments. Contacts between the myelin and axonal membranes involve multiple protein-protein, protein-lipid and possibly also lipid-lipid interactions (2–5). A key contact site, known as the paranode, is the region where the myelin sheath is anchored tightly to the axonal membrane, stabilised by the interaction between oligodendrocyte-specific and neurone-specific adhesion proteins (6). The paranode is also highly enriched in a specific class of lipids, the glycosphingolipids (GSLs), that partition into membrane microdomains in the outer leaflet of the plasma membrane and have numerous important roles in the function of the nervous system (7, 8). The importance of paranodal contacts is highlighted by the devastating demyelinating diseases that result from defects in the proteins that form these contacts or in changes to glycosphingolipid metabolism (9–11).

The galactose-based glycosphingolipids, including galactosylceramide (GalCer) and its 3-*O*-sulfated derivative sulfatide, are highly enriched in the myelin membrane and are important for myelin integrity (12–14). The sphingolipidoses Krabbe Disease and Metachromatic Leukodystrophy are fatal early-onset demyelinating diseases caused by defects in the enzymes β-galactosylceramidase (GALC) and arylsulfatase A (ARSA) that degrade GalCer and sulfatide, respectively (**Fig. 1A**) (15). These enzymes function in the lysosome to catalyse the stepwise removal of lipid headgroup moieties, facilitating the efficient recycling of these bioactive lipids. In disease, the lipid substrates and their deacylated derivatives, such as psychosine, accumulate and cause cell death. The majority of work to date has focussed on the deleterious effects of lipid accumulation on lysosomal function and on the detergent-like properties of psychosine (16–18). Substantial evidence supports the “psychosine hypothesis” for Krabbe disease pathology and blocking deacylation of GalCer to prevent psychosine production reduces Krabbe disease severity in mice (19). However, this work clearly demonstrated that psychosine is not the sole driver of disease and there is growing evidence in this and related sphingolipidoses that sphingolipid substrates accumulate throughout the cell, including at the plasma membrane (20–23). Sphingolipids also contribute to protein trafficking within the cell (24–26) and GSL imbalances change membrane protein abundance and function (23).

**Figure 1.**
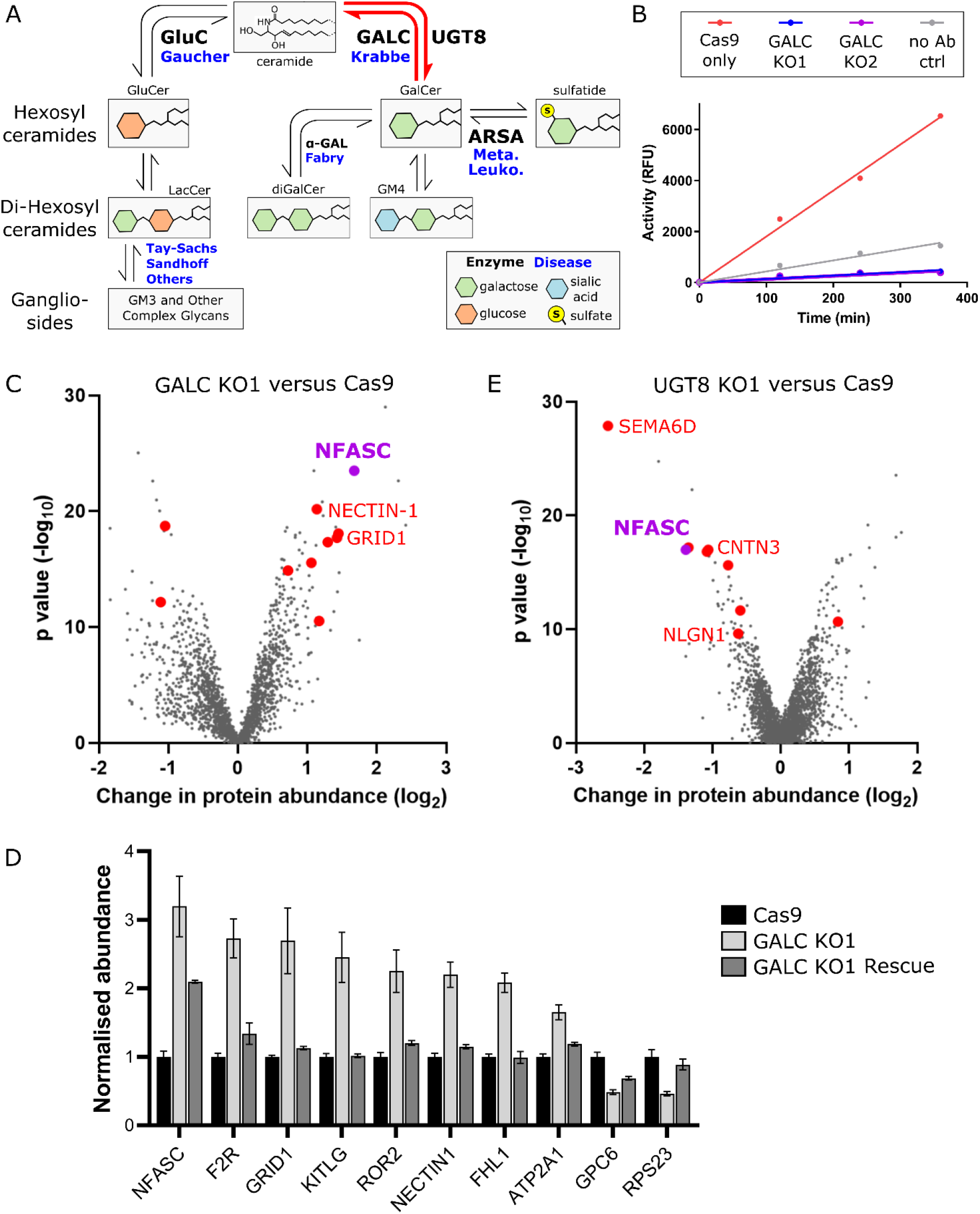
Disrupting galactosphingolipid metabolism alters the plasma membrane protein composition. **(A)** Simplified schematic diagram of GSL metabolism. Sequential addition or removal of glycans and sialic acid moieties produces a broad range of GSL headgroups. **(B)** GALC activity assays following immunoprecipitation (IP) from control and GALC knockout (KO) cell lines demonstrate loss of activity in the two clonal knockout lines compared with Cas9-only control cells and a no antibody (no Ab) control IP. Data are representative of n>3 independent experiments. **(C)** Quantitative mass spectrometry following enrichment of plasma membrane proteins (PMP-MS) from a GALC KO1 cell line compared with the Cas9 control. A volcano plot with the horizontal axis showing average fold change across three biological replicates and the vertical axis showing significance (two-sided t test) across the three replicates. The ten high-confidence targets (criteria as detailed in the main text) are coloured in red with select targets labelled. **(D)** Normalised protein abundance values, from the PMP-MS data, for high-confidence targets in the Cas9 control, GALC KO1 and GALC rescue cell lines. Normalised abundances have been used to allow comparison across proteins with different absolute abundances in the cell (mean ± SE, n = 3). Additional PMP-MS supporting data are available in Fig. S4A and Table S1. **(E)** As for panel (C) but for a UGT8 KO cell line. Additional PMP-MS supporting data are available in Fig. S4B and Table S2.

Mouse models of Krabbe disease and mice where GSL-synthetic enzymes have been ablated all display hypomyelination involving specific destabilisation of contacts at the paranodal junction (13, 27–31). These models demonstrate that it is not just accumulation of GSLs, but also their absence, via loss of the UDP-galactosyl transferase 8 (UGT8 a.k.a. ceramide galactosyl-transferase, CGT) or ceramide sulfotransferase (CST) that can cause severe and rapid demyelination. As loss of GSL synthetic enzymes would not be expected to result in increased psychosine abundance, the molecular mechanisms underlying demyelination in these diseases may be more complex than psychosine-mediated cell death, and involve specific changes at myelin-axon contact sites. We sought to determine whether the plasma membrane protein composition is changed in cell-based models of sphingolipid disorders and how these changes may contribute to disease phenotypes.

Here we exploit oligodendrocyte-like cells to show that disruption of galactosphingolipid metabolism via deletion of the enzymes GALC and UGT8 specifically changes the plasma membrane proteome. The cell adhesion protein Neurofascin (NFASC) is particularly sensitive to GSL imbalances, being reciprocally altered in its abundance at the PM when GSL synthesis and catabolism is inhibited. We demonstrate direct binding of the oligodendrocyte isoform of NFASC, NF155, to sulfatide but not to other closely related galactosphingolipids. Mapping of NF155 binding to sulfatide and structural characterisation of its sulfatide-binding extracellular domain shows how this essential component of the paranode may be oriented relative to the myelin membrane, enabling the tight membrane apposition required for paranode stability.

## RESULTS

### Disrupting galactosphingolipid metabolism alters the plasma membrane protein composition

Krabbe disease pathology occurs in patients who possess <10% activity of the catalytic enzyme GALC. The majority of Krabbe disease patients possess a 30 kb deletion within this gene in at least one allele (18). Therefore, CRISPR/Cas9-mediated deletion of GALC represents a good model for this disease. Multiple independent clonal knockouts of GALC were produced in the oligodendrocyte-like cell line, MO3.13 (32, 33). Genomic editing within early exons of the GALC gene was verified by sequencing across the guide RNA (gRNA)-target sites (**Fig. S1A** and **B**). To test for GALC enzyme activity in these cells, a highly sensitive activity assay was performed following enrichment of any residual GALC protein via immunoprecipitation (**Fig. 1B**). These data confirm loss of GALC activity in two independent GALC KO cell lines generated using gRNAs targeting different regions of the gene. To facilitate comparative analysis of how galactose-based GSLs influence cell function, additional cell lines were generated knocking out the synthetic enzyme UGT8. This enzyme functions in the ER to add the galactose headgroup to ceramide, forming the backbone of all galactosylated-sphingolipids (**Fig. 1A**). Gene editing was verified by sequencing across the gRNA-target sites (**Fig. S1C** and **D**). Mass spectrometry-based lipidomic analysis specifically tailored for detection of glycosylated sphingolipids demonstrated accumulation of several GSL species in the GALC KO cells and reduced abundance of GSLs in the UGT8 KO cells (**Fig. S2**).

We quantified the differential expression of plasma membrane proteins between wild-type (WT) cells, Cas9-only control cells, the two independent GALC KO clones generated using different gRNAs and a GALC rescue line where GALC expression had been restored **(Fig. S3)**. Intact cell monolayers were labelled with activated aminoxybiotin to allow enrichment of surface proteins via streptavidin-affinity prior to mass spectrometry analysis (34). Plasma membrane profiling mass spectrometry (PMP-MS) data were collected for 3460 proteins quantified in all datasets allowing several criteria to be used for determining confidence in the target identification. Protein abundance changes in the two independent GALC KO cell lines were compared with Cas9 controls (**Fig. 1C** and **Fig. S4A**) and high-confidence targets were selected based on: no significant change between WT and Cas9-only control lines; significant change in membrane abundance in both knockout lines; change in abundance by more than 2-fold in at least one KO line; identification by two or more unique peptides, and significant recovery following re-expression of GALC (**Fig. 1D**). These stringent criteria identified ten high-confidence hits, eight of which showed increased abundance at the cell surface of KO cells (**Table S1**). These proteins include NECTIN-1, NFASC and GRID1 (a.k.a. GluD1) with important roles in cell adhesion, axon-glial contacts, synapse formation and neurotransmission (35–37), highlighting their potential roles in the demyelinating neurodegenerative pathologies seen in Krabbe disease. As only a small subset of PM proteins were altered in these cell lines, the observed changes are likely to be caused by direct effects rather than by non-specific fluctuations in membrane fluidity that would be expected to influence the abundance of many PM proteins indirectly.

To probe which of the GALC high-confidence hits may be directly sensitive to changes in GSL abundance, as opposed to downstream cellular consequences of these changes, PMP-MS analysis was also performed on the UGT8 KO cell lines and compared to Cas9 control cells, applying equivalent criteria as above (**Fig. 1E**and **Fig. S4B**). Nine high-confidence hits were significantly altered in both UGT8 KO cell lines (**Table S2**). All but one of these proteins has lower abundance at the PM versus the control cells, in a pattern opposite to that seen for the GALC KO cells. The high-confidence hits again include several proteins, such as CNTN3, NLGN1 and SEMA6D with important roles in cell adhesion and neuronal function (38–40). Interestingly, NFASC (Neurofascin) was identified again as a high-confidence target with reciprocal changes in cell surface abundance, being increased in GALC KO cells and decreased in UGT8 KOs (**Fig. 1C** and **E**). NFASC is an L1-type cell adhesion protein from the immunoglobulin superfamily, with crucial roles in neuronal development and myelination (36). Loss of a specific isoform of NFASC (NF155) leads to disorganisation of the paranode (41) and causes a severe neurodevelopmental disorder (42). NF155 is part of a multi-protein complex at the paranode that bridges the myelin-axon contact site (43, 44) and the stability of this complex depends on the presence of GSLs (3, 45). Interestingly, the PM proteomic data determined here for both GALC and UGT8 KOs identify that NFASC abundance at the cell surface is sensitive to, and correlated with, GSL abundance.

### The NFASC isoform NF155 specifically and directly binds sulfatide

NFASC possesses multiple splice variants, each with distinct functions, localisation and expression patterns (46). To probe the mechanism underlying NFASC abundance changes in response to sphingolipid imbalances, we focussed on the two most well-studied isoforms expressed in the CNS, NF155 and NF186 (**Fig. 2A**). NF155 is expressed specifically in oligodendrocytes and localises to the paranode, where it forms a tight septate-like junction between the myelin and axonal membranes through interaction with the axonal proteins Contactin 1 (CNTN1) and Contactin associated protein (Caspr) (43, 47). This junction is crucial as a diffusion barrier between the voltage-gated sodium channels (NaVs) in the node and voltage-gated potassium channels in the juxtaparanode. In contrast, NF186 is expressed in neurones and localises to the nodes of Ranvier, where it clusters NaVs and interacts with components of the extracellular matrix, contributing to long-term node stability (48). Therefore, both isoforms play key but distinct roles in promoting efficient signal conduction along the neuronal axon. These two isoforms differ in domain organisation, particularly at the C-terminal end of the extracellular domain (ECD) where NF155 has four fibronectin type-III (Fn) domains (Fn1-4), while NF186 has Fn domains 1, 2, 4 and 5 with a PAT (**P**roline/**A**lanine/**T**hreonine, mucin-like) domain between Fn4 and 5 (**Fig. 2A and Fig. S5**). To test whether NFASC isoforms bind GSLs directly, and to probe their binding specificity, the full-length ECDs of NF155 and NF186 were expressed in the human cell line HEK293-F, secreted into the medium and purified maintaining native glycans for testing in liposome interaction assays. Three galactosylated sphingolipids: GalCer, sulfatide and GM4, were individually incorporated into liposomes containing phosphotidylcholine (PC) and rhodamine B-labelled phosphotidylethylolamine (Rhod-PE) and tested for binding to full-length NF155-ECD and NF186-ECD (**Fig. 2B**). Both NF155 and NF186 possess some binding to PC-containing liposomes. However, despite NF186 and NF155 being closely related isoforms, only NF155 displayed dramatically increased binding to liposomes containing sulfatide with no increased binding to liposomes containing the closely related lipids GalCer or GM4. To ensure that this binding was specific for sulfatide and not mediated by non-specific electrostatic interactions, an alternative anionic lipid, phosphatidylserine (PS) was tested in liposome binding assays with no detectable binding (**Fig. S6**).

**Figure 2.**
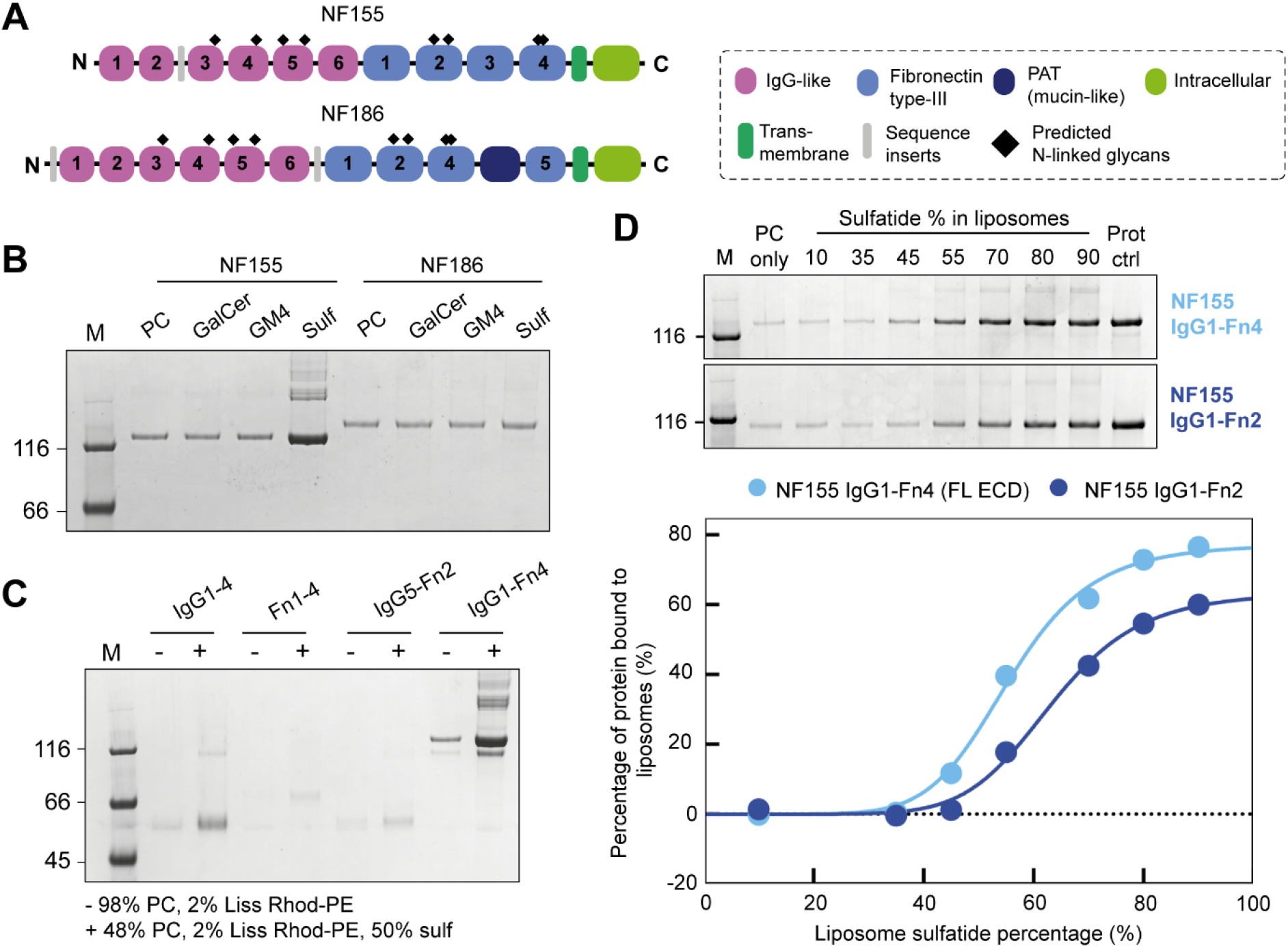
Lipid-binding assays demonstrate that NF155 specifically and directly binds sulfatide. **(A)** Protein domain schematic of NF155 and NF186, highlighting the differences between the two, arising from alternative splicing. The IgG-like domains are in pink, the Fn3-like domains are in light blue, the PAT domain is in dark blue, the transmembrane domain is in dark green, the intracellular domain is in light green, and the predicted N-glycosylation sites are highlighted by black diamonds. **(B)** Liposome binding assay demonstrating that NF155-ECD binds specifically to liposomes containing sulfatide, and not to the other closely-related GSLs, GalCer and GM4, while NF186 does not bind (above background). Assays were performed with 1 μM protein and 0.8 mM liposomes. PC liposomes contain 98% PC and 2% Rhod-PE. GalCer, GM4 and Sulf liposomes contain 48% PC, 2% Rhod-PE and 50% of either GalCer, GM4 or sulfatide, respectively. **(C)** Mapping the sulfatide binding region using NF155-ECD truncations (1 μM protein, 0.8 mM liposomes) demonstrates that only the full-length (FL) ECD shows significant binding to liposomes containing 50% sulfatide. **(D)** *Top:* Liposome binding assay performed with 250 nM NF155 using 1.6 mM liposomes containing increasing concentrations of sulfatide (IgG1-Fn4, top gel, IgG1-Fn2, lower gel). *Bottom:* Densitometric analysis of the SDS PAGE reveals a sigmoidal binding relationship for both NF155 constructs (IgG1-Fn4, light blue, IgG1-Fn2, dark blue). The data shown is the quantification are from the gels shown above and are representative of four independent experiments. Prot ctrl represents total protein added to each assay sample. Replicate data are provided in Fig. S8. A total of n=4 independent experiments were performed.

To explore the potential effects of NF155 glycosylation on sulfatide binding, we produced NF155-ECD in the presence of kifunensine, to yield protein with high-mannose glycosylation, which can be removed with endoglycosidase H (Endo H). The high-mannose and deglycosylated forms of NF155-ECD showed the same binding to sulfatide-containing liposomes as the standard complex human glycosylated form (**Fig. S7**), supporting that the interaction with sulfatide is mediated by direct binding to the protein and is not a glycan-glycan interaction.

To map which domain (s) mediate sulfatide binding, NF155 truncations were expressed and purified encompassing regions that differ between NF155 and NF186 isoforms: the four N-terminal Ig domains (IgG1-4); the four C-terminal Fn domains (Fn1-4); and a central region (IgG5-Fn2). Liposomes containing 50% sulfatide were used for interaction assays, revealing that IgG1-4 retains some sulfatide binding, while Fn1-4 and IgG5-Fn2 possess barely detectable binding to liposomes (**Fig. 2C**). Interestingly, the full-length NF155-ECD shows much stronger binding to sulfatide-containing liposomes than any truncation, suggesting avidity-enhanced binding to multiple, distinct and weak sulfatide-binding sites. To assess potential binding co-operativity, binding assays were performed using 1.6 mM liposomes containing increasing concentrations of sulfatide (0-90%) and incubated with 250 nM NF155-ECD (**Fig. 2D** and **Fig. S8**, *light blue*). Densitometric quantification of NF155 bands following SDS-PAGE identified a sigmoidal relationship between NF155 binding and the sulfatide content of liposomes (**Fig. 2D**). Fitting the data to the Hill equation gives a Hill coefficient (7.7 ± 0.9, mean ± SE) consistent with NF155 possessing multiple sulfatide binding sites and that the binding of ligands is co-operative.

Domain mapping experiments suggested that the membrane-proximal C-terminal Fn domains are dispensable for the interaction with sulfatide (**Fig. 2C**). Therefore, a construct of the NF155-ECD missing Fn3-4 (IgG1-Fn2) was purified and assayed for binding to sulfatide-containing liposomes (0-90%) (**Fig. 2D** and **Fig. S8**, *dark blue*). Surprisingly, IgG1-Fn2 showed a reduction in sulfatide binding when compared with the full-length ECD, which is not caused by protein aggregation or oligermisation as a consequence of the truncation as shown by multi-angle light scattering coupled to size-exclusion chromatography (SEC-MALS) analysis (**Fig. S9**). The fraction of protein that binds to liposomes is significantly decreased (IgG1-Fn2: 55.0% ± 7.0%, IgG1-Fn4: 76.8% ± 2.1% at saturating sulfatide concentrations; mean ± SE, n = 4, paired t test p = 0.025), and the percentage sulfatide required for half-maximal binding was significantly increased (IgG1-Fn2: 62.3% ± 2.1%, IgG1-Fn4: 55.8% ± 1.4%; p = 0.02) suggesting increased membrane dissociation for the truncation. However the Hill coefficient was not significantly different (IgG1-Fn2: 8.4 ± 0.7; IgG1-Fn4: 7.7 ± 0.9; p = 0.3). This suggests that removal of the last two Fn domains did not remove a sulfatide binding site but the presence of these domains increases the affinity of NF155 for sulfatide. The conformation of the full-length ECD may therefore be important for efficient membrane association.

### NF155-ECD adopts an S-shape

To date, the only experimental structural data available for NFASC is a crystal structure of the four N-terminal Ig domains (49) that fold back upon themselves to form a globular “horseshoe” arrangement. This work also suggested that NFASC may form homodimers (49). To explore how the conformation of the full-length NF155-ECD may contribute to its membrane binding behaviour we employed a range of biophysical and structural techniques. Using SEC-MALS we demonstrated that the full-length, glycosylated NF155-ECD is a monomer (**Fig. 3A**). This difference with the published literature may be attributed to previous studies using truncated NFASC constructs or the removal of surface glycans, as we observed that deglycosylation of NF155 destabilises the protein resulting in time-dependent aggregation and degradation. As our data have implicated that the full-length ECD is required for binding, subsequent structural studies were carried out using the fully glycosylated full-length ECD. Attempts to crystallise this construct were unsuccessful, potentially due to the heterogeneity of the glycans. Recent advances in artificial intelligence-based deep learning strategies, exemplified by AlphaFold2 (AF2), allow for the highly accurate prediction of protein structures (50). Structural predictions of NF155 using ColabFold (51), implementing AF2, provide accurate models for the individual domains of NFASC. However, the overall conformation and relative inter-domain orientations (described by the Predicted Alignment Error (PAE) plot) are poorly predicted outside the IgG1-4 region, limiting our understanding of the overall ECD shape (**Fig. S10**). The NF155-ECD is near the lower size limit for cryo-electron microscopy (EM) structure determination and preliminary data collection did not yield high quality particle analysis. We therefore used negative stain EM to capture images of the overall ECD structure. Raw images of the NF155 full-length ECD sample reveal clear, high-contrast particles that appear monodisperse and homogenous (**Fig. 3B**). In total, 26,642 particles were analysed, allowing class averaging (**Fig. 3C**) and single-particle reconstruction (**Fig. 3D**). This three-dimensional reconstruction, at 19.2 Å (**Fig. S11)**, reveals that the NF155-ECD possesses an asymmetric S-shaped conformation with a bulbous “head” followed by two sharp bends (**Fig. 3D**). This structure also shows that, although the first four IgG domains are known to fold back upon themselves, the full ECD is not globular, adopting a relatively flat shape with dimensions 19 × 11 × 6 nm (**Fig. 3D**).

**Figure 3.**
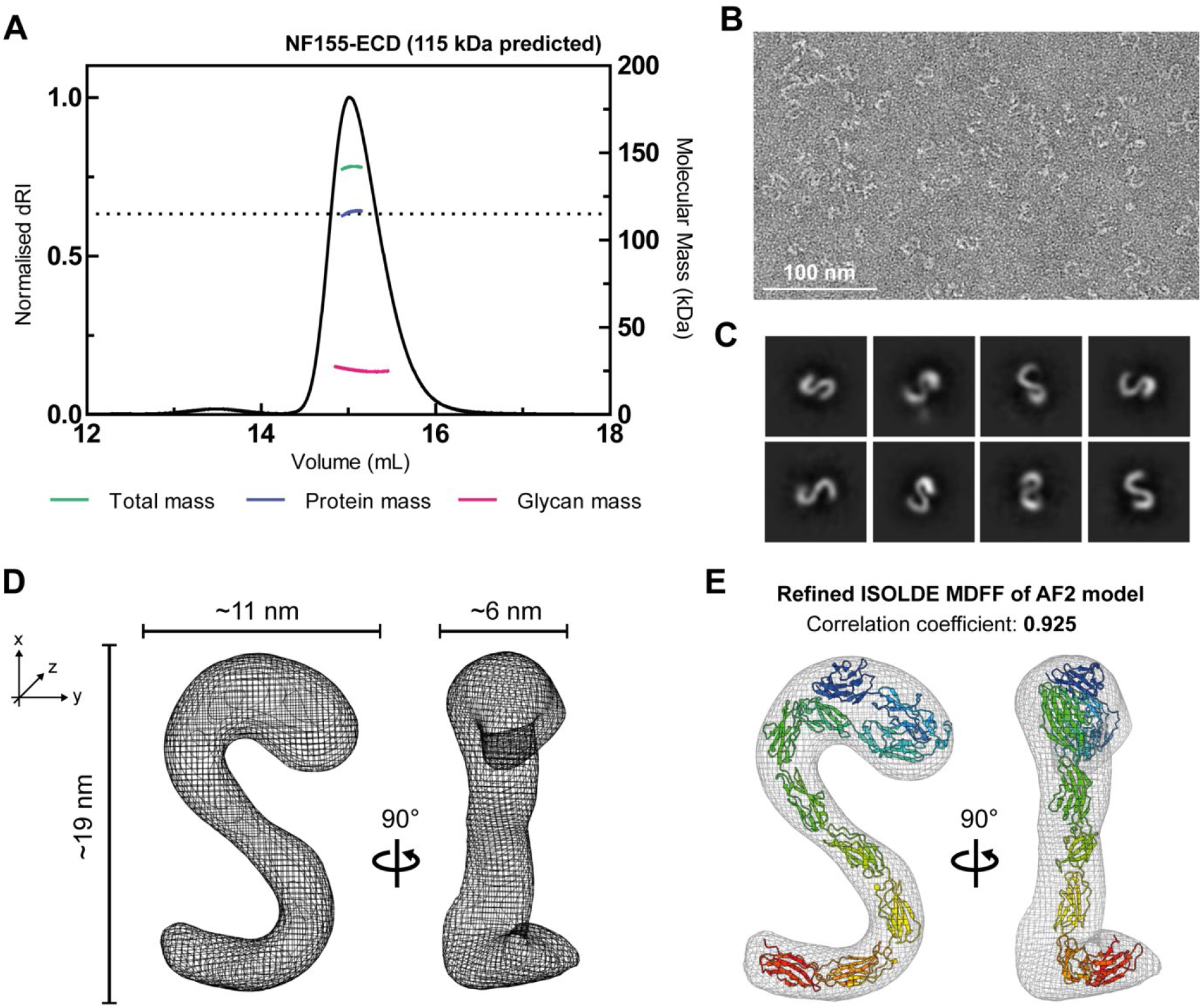
The extracellular domain (ECD) of NF155 is monomeric and adopts an “S”-shape. **(A)** Size exclusion chromatography coupled to multi-angle light scattering (SEC-MALS) demonstrates that the NF155 FL ECD is monomeric with a protein mass of 115 kDa (consistent with the predicted mass, dotted line) and 25 kDa glycans. **(B)** Raw image of NF155 FL ECD single particles, stained with 2% uranyl acetate. **(C)** Selection of 2D class averages generated from the negative stain EM data using CryoSPARC. **(D)** The 3D reconstruction of the NF155 FL ECD, generated using CryoSPARC analysis of the negative stain EM data. The particle possesses dimensions 19 x 11 x 6 nm, two orientations are shown rotated by 90°. **(E)** The best AF2-generated model was fit to the EM map using ISOLDE molecular dynamics flexible fitting (MDFF), resulting in a high quality fit (correlation coefficient = 0.925).

Initial rigid-body docking of the AF2 models for the full-length NF155-ECD did not fit the EM density maps well (**Fig. S12A**), with modest correlation coefficients (CC = 0.748-0.859). Flexible fitting of the best AF2 model (model rank 1) into the EM map using molecular dynamics simulations as implemented by ISOLDE (52) in ChimeraX (53) allowed for a better fit (CC = 0.913, **Fig. S12B**). However, visual inspection of the fitted structure into the EM map suggested that the “head”domain encompassing IgG1-4 was not oriented optimally to explain the density. Therefore, the IgG1-4 domain was manually rotated as a rigid body, maintaining connectivity to IgG5. Following molecular dynamics based flexible fitting, as above, the final model displayed an excellent fit to the map (CC = 0.925, **Fig. 3E**).

### NF155 membrane binding orientation

NF155 is localised to the paranode, a septate-like junction between myelin and axonal membranes that are separated by a gap of around 3-8 nm (54, 55). The dimensions of the NF155-ECD structure are 19 × 11 × 6 nm (**Fig. 3C**), suggesting that in order to fit within this tight inter-membrane space, NF155 must lie flat (**Fig. 4A**). As both NF155 and sulfatide reside in the membrane of myelin-forming cells, the sulfatide-binding data suggest that NF155 binds in *cis* to the membrane within which it is embedded. To test this hypothesis, a liposome clustering assay was developed incorporating different fluorescent lipids to facilitate direct visualisation of the interaction. At high concentrations NF155 caused liposomes to coalesce forming large assemblies that compromise visualisation (**Fig. S13A**). Therefore, these assays employed a relatively low concentration of NF155 (20 nM). For initial clustering assays, one population of liposomes was produced incorporating Rhod-PE (pink) and sulfatide while a second population contained NBD-PE (green) and Ni-NTA conjugated lipids (DGS-Ni-NTA). Following mixing, liposomes were inspected by wide-field microscopy and did not show clustering of different coloured liposome populations (**Fig. 4Bi**). Upon addition of NF155-ECD, which can be incorporated via its C-terminal His_6_ tag into the green Ni-containing liposomes, significant clustering of pink and green liposomes is observed (**Fig. 4Bii** and **4C**). In this case, NF155-ECD clusters the different liposome populations by being anchored into the green Ni-containing liposomes and binding to the sulfatide present in the pink liposomes (**Fig. 4B**, schematic). Additional clustering assays were carried out to test if incorporation of sulfatide into both liposomes could inhibit clustering (**Fig. 4Biii** and **iv**). Incorporation of sulfatide into both liposomes did inhibit clustering in the presence of NF155-ECD suggesting that NF155 can bind to the membrane within which it is embedded (binding in *cis*) and that this buries the sulfatide-binding sites. Taken together, these data reveal an intriguing membrane association model for NF155, in which the ECD has an extended sulfatide-dependent interaction with the membrane it is embedded in, maintaining it in a “flat” conformation relative to the membrane that is consistent with the size of the intermembrane space at the paranode.

**Figure 4.**
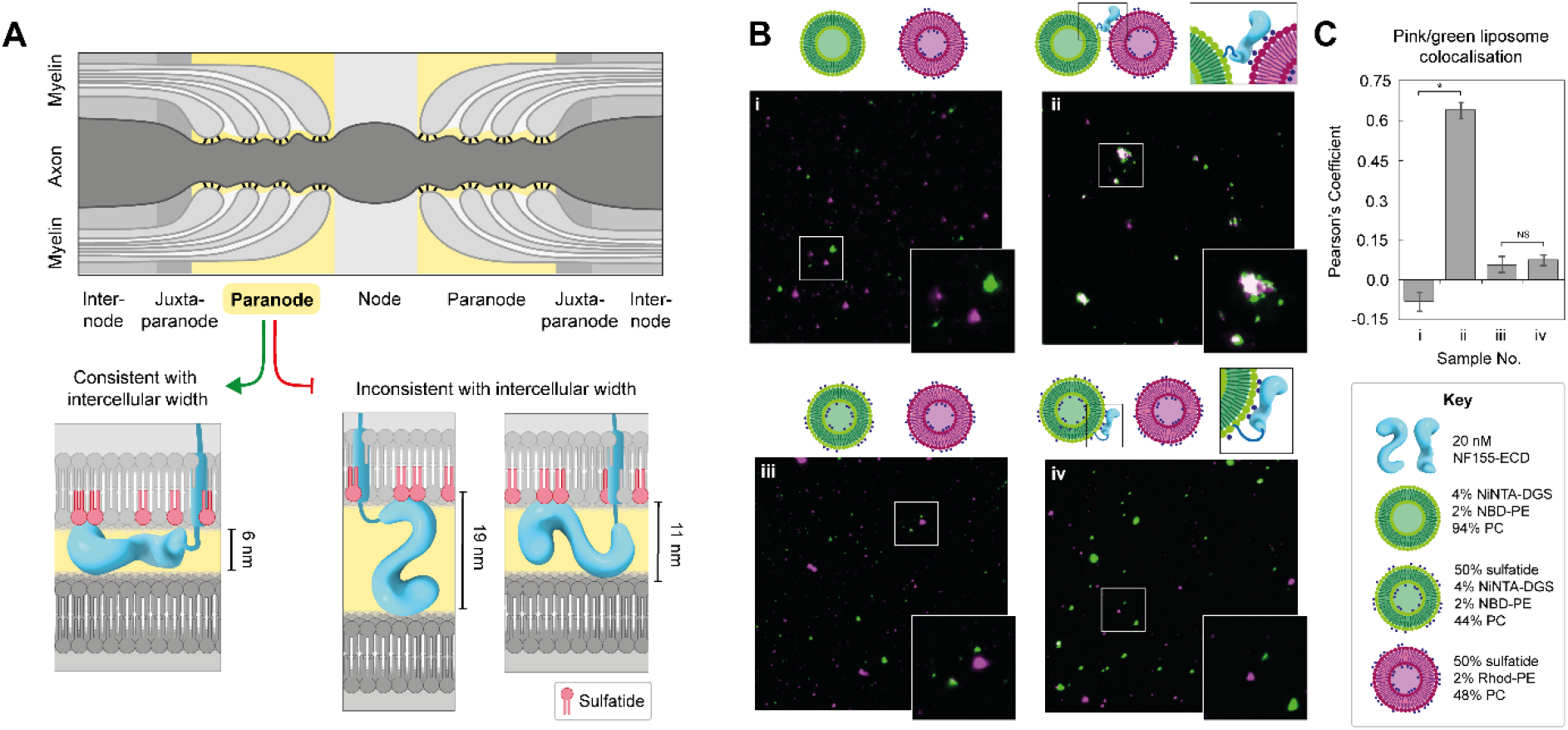
NF155 membrane orientation. **(A)** The paranodal junction (yellow) is estimated to possess an intermembrane gap of 3-8 nm, a width that is only consistent with a model of NF155 FL ECD lying flat along the myelin membrane, potentially stabilised by its interaction with multiple sulfatide headgroups (red). **(B)** Liposome binding assays performed at 20 nM NF155 with 0.4 mM liposomes, demonstrating clustering of NBD-labelled liposomes containing 1% NiNTA-DGS (green) with Rhodamine-labelled liposomes containing 50% sulfatide (pink) in the presence of the NF155-ECD (sample ii). This clustering is prevented when NF155 is preincubated with NBD-labelled liposomes containing 50% sulfatide (sample iv). No clustering is seen when the liposomes are incubated in the absence of NF155-ECD (samples i and iii). Images shown here represent 8% of the full image area. **(C)** Quantification of the colocalization of the pink and green liposomes in the samples described in (B). Mean ± SD is shown for quantification from 3 full images. Significance calculated using a two-tailed unpaired t-test, *p ≤ 0.0001, NS = not significant.

## DISCUSSION

Here we have demonstrated that disruption of GSL metabolism in oligodendrocyte-like cells alters the abundance of PM proteins. Loss of the GSL metabolic enzymes GALC and UGT8, which remove or add a galactose headgroup from/to ceramide, respectively, results in significant changes in the abundance of specific plasma membrane proteins. Importantly, the small subset of altered proteins suggests that these changes are direct effects mediated by changes in the abundance of specific lipids arising from the loss of these enzymes, rather than non-specific modulation to membrane fluidity. Of particular interest is the observation that NFASC shows reciprocal changes in abundance that correlate with GSL abundance. This finding suggested that a direct NFASC-GSL interaction was driving the change in NFASC abundance at the PM. Here we have directly demonstrated *in vitro* that the NF155 isoform of NFASC interacts specifically with sulfatide but not with other closely related or similarly charged lipid species, while the NF186 isoform does not. Despite being extremely similar (**Fig. 2A** and **S5**), these two isoforms have different roles within the nervous system that, as demonstrated by our data, are likely driven by distinct interactions with other cellular components. Direct binding of the NF155-ECD to sulfatide demonstrated here provides a molecular explanation for previous observations that NF155, but not NF186, is resistant to detergent solubilisation, suggesting it associates with lipid rafts *in vivo* (56) and that this requires sulfatide (57).

The PMP-MS data for the GALC rescue cell line provides strong evidence for the direct importance of GALC in maintaining PM homeostasis. The restoration of protein abundances back to normal levels was excellent for the ten high-confidence targets (**Fig. 1D**), confirming that these were direct effects of the loss of GALC and not off-target effects. Despite significant recovery, the abundance of NFASC at the PM recovered less well than other targets. Re-expression of GALC was induced by addition of doxycycline prior to differentiation for 7 days, and this may be insufficient to restore key GSL species to normal levels. The direct binding of NFASC to sulfatide may make it especially sensitive to cellular lipid composition, requiring a longer rescue of GALC expression for full recovery. The data presented here demonstrating reciprocal changes in NFASC abundance at the PM in cells accumulating (GALC KO) or deficient (UGT8 KO) in GSLs, along with the demonstration of direct binding of NF155 to sulfatide, suggests several potential mechanisms for how NFASC accumulates in membranes. Direct binding supports the hypothesis that NFASC and sulfatide traffic together through the secretory and endosomal pathways, resulting in a change in abundance for both in different cellular compartments, including the PM. Direct binding may also result in increased stability of NF155, reducing its endocytosis, or proteolytic shedding (58) from the PM. However, the precise mechanism of NFASC accumulation at the PM in response to GSL changes remains unclear.

Liposome binding assays were conducted using truncations of NF155 to map the specific sulfatide-binding domain(s). Both N- and C-terminal truncations significantly reduced binding to sulfatide-rich membranes and the binding of the full-length NF155-ECD revealed that it possesses multiple sulfatide binding sites that display positive cooperativity. Due to the relatively small size of the sulfatide headgroup, a single galactose with one sulfate group, this avidity effect would help facilitate high affinity binding to sulfatide-containing membranes. Sulfatide is highly enriched at myelin-axon contact sites, as demonstrated by anti-sulfatide antibody binding to myelinated axons (59). This enrichment may result in extremely high local concentrations of sulfatide, helping facilitate multi-valent interactions. Interestingly, these data do not support traditional representations of NF155 as a linear arrangement of tandem Ig and Fn domains extending away from the PM. Instead, for NF155 to engage with several sulfatide headgroups along its length it must either fold back towards or lie flat along the membrane. Our structural analysis shows that the full-length NF155-ECD adopts an S-shaped conformation, bending back upon itself several times, with a dense N-terminal “head” domain and a highly curved C-terminal region. Bent or curved structural conformations, such as seen here for the NF155-ECD, have been observed previously for other proteins involved in cell-cell contacts such as contactins and sidekick proteins, where their shape was proposed to facilitate both *cis* and *trans* interactions with membranes or other membrane proteins (60, 61). In these cases, this arrangement was inferred from structures of fragments of these proteins rather than full-length ECD conformations. Importantly, the dimensions of the NF155-ECD structure combined with liposome clustering assays support that the ECD of NF155 lies flat within the gap at the paranode, contacting the myelin membrane. This extensive membrane interaction is consistent with our observations that NF155 binds sulfatide via multiple sites along the ECD. In a CST KO mouse model, loss of NF155 at the paranode was age dependent, suggesting sulfatide is not necessary for initial formation of the paranodal junction but very important for its long-term stability. Based on our data this stability may be driven by the multiple sulfatide binding sites in NF155 forming an extensive interaction with the membrane that could promote tightening of the paranodal gap and facilitate interactions with axonal proteins, CNTN1 and Caspr, by displaying the correct interaction interfaces.

Previous work using mouse models where galactose-based sphingolipids are dysregulated by either accumulation due to GALC loss (Twitcher mouse), or deficiency due to UGT8 or CST loss, all manifest with defective myelination (13, 28, 31). Interestingly, this means that rather than reciprocal changes in lipid abundance leading to reciprocal changes in myelin integrity, it is the precise homeostasis of GSL levels that is essential for stable myelin formation. Lipidomics data from GALC and UGT8 KO cell lines suggest that disruption to GSL metabolism results in complex lipid changes, including altered abundance of several GSL species (**Fig. S2**). This is consistent with previous studies showing unexpected disruption to sphingomyelin and phospholipid abundances in CST KO mice (62), supporting that it is not just the direct enzyme substrates that accumulate in GSL diseases. This complexity makes it challenging to determine which lipid species may be the primary driver of myelination defects and it seems likely that a combination of changes to multiple lipid species and several membrane proteins drives severe disease. Although sulfatides were not able to be quantified in these cell lines, the changes to NFASC abundance are consistent with changes seen in models and diseases of sulfatide imbalance. Specifically, CST KO mice display node/paranode disorganisation and suffer from age-dependent neurodegeneration due to loss of paranodal NF155, while NF186 localisation to the node is unchanged (3). Similarly, patients with Metachromatic Leukodystrophy are unable to break down sulfatide due to the loss of ARSA, leading to sulfatide accumulation in both the lysosome and myelin membrane, causing rapid demyelination (23). These data support that both accumulation and absence of sulfatide causes myelination defects, and that NF155 and sulfatide may play unexpected but important roles in the pathology of Krabbe disease.

Understanding how loss of NF155, a crucial junctional protein, would destabilise axon-myelin contacts is clear, but it is less obvious how increased abundance might also destabilise these contacts. It is possible that accumulation of NF155 at the PM of myelin-forming cells might interfere with the correct formation of the axon-myelin protein complexes, specifically the interactions between NF155 and the axonal proteins CNTN1 and Caspr. Alternatively, the high abundance of NF155 combined with its ability to interact with membranes may have a destabilising effect on the membrane similar to that caused by membrane-embedding antimicrobial peptides (63). In support of this, we observed that at high concentrations, NF155 can induce membrane destabilisation and blebbing of liposomes and cells *in vitro* (**Fig. S13B**). However, the relevance of this to demyelinating processes *in vivo* remains unclear. Finally, there is evidence that during myelin compaction there must be reduction in the charged components of the glycocalyx, including sialic acid residues, in order to avoid electrostatic repulsion between membranes (64). It is therefore possible that, in addition to a specific role for NF155 in Krabbe disease, increased abundance of other lipids and proteins, as seen in the GALC KO, may hinder myelination via increased electrostatic repulsion.

This work demonstrates the exquisite sensitivity of the PM proteome to alterations in lipid homeostasis. Several PM proteins with important roles in cell adhesion, neurotransmission and synapse formation were altered in this cell-based model of Krabbe disease, highlighting potential new leads in understanding the molecular consequences of GSL accumulation. Our work focussed on NFASC, directly demonstrating its specific and highly cooperative GSL binding activity and revealing the first full-length ECD structure of a paranodal complex protein. Taken together, these results suggest that the NF155-ECD interacts extensively with the myelin membrane in which it is embedded, in a manner distinct from the classical paradigm. Future work investigating the roles of other axonal proteins and complex GSLs within the paranodal junction will shed more light on the molecular details of myelin stabilisation, leading to a more comprehensive understanding of the pathophysiology of demyelinating diseases.

## METHODS

### Cells and cell culture

Human oligodendrocyte-like MO3.13 cells were a gift from Prof. Louise Boyle (University of Cambridge). For normal maintenance, cells were cultured in DMEM containing 4500 mg/L-glucose, L-glutamine, sodium pyruvate, and sodium bicarbonate supplemented with 10% (v/v) foetal calf serum (FCS). Differentiation was induced by culturing in DMEM, 1% (v/v) FCS and 100 nM phorbol 12-myristate-13-acetate (PMA) for 7 days with fresh medium applied every 2-3 days. Cells were cultured at 37°C in a humidified 5% CO_2_ atmosphere.

### CRISPR/Cas9 genome editing

Oligonucleotides encoding single guide RNAs (sgRNAs, **Table S3**) targeting human GALC exon 3 and UGT8 exon 2 were cloned into the *BbsI* restriction site in pSpCas9(BB)-2A-Puro as previously described (65). MO3.13 cells were transfected with 2 μg of each plasmid (including an untargeted Cas9 control) using FuGeneHD transfection reagent as per manufacturer’s instructions. Following overnight incubation, selection with 2 μg/mL puromycin was applied for 2 days. Clonal cell lines were established by limiting dilution. After expansion of clones, editing at the relevant genomic site was confirmed by amplification and sequencing of PCR product from genomic DNA (**Table S4**) and analysed using TIDE (66).

### Generation of GALC rescue cell line

Human GALC cDNA was cloned into the lentiviral vector pCW57 using restriction sites *Nhe*I and *Sal*I. For lentiviral production 6 μg of pCW57-hGALC was co-transfected into 1.2 x 10^7^ HEK293-T cells with 12 μg psPAX2 and 3 μg pMD2.G viral packaging and envelope vectors (gifts from Dr Hayley Sharpe, Babraham Institute, Cambridge) using FuGeneHD transfection reagent. After 8 hrs, transfection medium was replaced with fresh DMEM containing 10% (v/v) FCS and incubated for 48 hrs. Virus-containing supernatant was concentrated 200-fold by ultracentrifugation (100,000 *g*, 1.5 hrs) and 25 μL was added dropwise to a 6-well plate containing MO3.13 GALC KO1 cells and incubated for 48 hrs. Following this, transduced cells were selected by incubation for 48 hrs in medium containing 2 μg/mL puromycin. The pCW57 vector allows for addition of doxycycline to induce expression of GALC. Doxycycline titrations were used in combination with GALC activity assays (see below and **Fig. S3**) to determine that 0.2 μg/mL doxycycline induced WT levels of GALC activity.

### Activity Assays

GALC activity assays were modified from those described previously (67). MO3.13 cells were lysed in 10 mM Tris pH 8.0, 1 mM EDTA, 1% Triton X100, 140 mM NaCl, 1 mM PMSF and pre-cleared by incubation with Protein A resin. Endogenous GALC protein was immunoprecipitated (IP) from cleared lysate using our monoclonal GALC antibodies described previously (68). Lysates were incubated at 4°C overnight with GALC antibody and Protein A resin. A control IP was carried out with Protein A resin but with no GALC antibody. Beads were washed four times with lysis buffer before being transferred into activity assay buffer (20mM sodium acetate, pH 4.6, 150 mM NaCl, 0.1% (v/v) NP40 containing fluorogenic substrate 0.3 mM 4-methylumbelliferyl β-D-galactopyranoside) pre-equilibrated to 37°C. The reaction product was monitored at three timepoints over six hours by fluorescence detection (excitation at 365 nm, emission at 445 nm) following addition of stopping buffer (480 mM NaOH, 380 mM glycine, pH 10.6).

### Lipidomics

For lipidomic analysis, 6 x 10^5^ MO3.13 cells were seeded into 15 cm dishes and incubated for 24 hrs in DMEM containing 10% (v/v) FCS. Medium was then replaced with differentiation medium (as described above) and incubated for a further 7 days. Cells were washed twice in ice-cold PBS and scraped into microfuge tubes. Cells were gently centrifuged (500 *g*, 5 mins, 4°C) and residual PBS removed. Pellets were snap frozen in liquid nitrogen and stored at −80°C. Lipids were extracted according to the modified Folch protocol (69). Cell pellets (25-28 mg) were spiked with 10 μL of Internal Standard mixture (**Table S5**). Two mL of chloroform and 1 mL of MeOH were added. The resulting mixture was homogenised in an ultrasonic bath for 15 min at 40°C. When samples reached ambient temperature, 600 μL of 250 mM of ammonium carbonate in water was added, and the samples were homogenised in the ultrasonic bath at 40°C for 15 min, then centrifuged at 3000 rpm for 5 min, and the organic layer (bottom layer) was transferred using Pasteur pipettes to 8 mL glass vials. The extraction was repeated by adding 2 mL of chloroform to the aqueous phase. Then, the samples were sonicated in the ultrasonic bath at 40°C for 15 min, centrifuged at 3000 rpm for 5 min, and finally, the lower organic layer was transferred and combined with the first layer. The combined organic phases were evaporated under the nitrogen stream to dryness, and the extracts were stored at −80°C.

Prior to analysis, samples were dissolved in 250 μL of chloroform/MeOH (1:1, v/v) mixture. An Agilent 1290 Infinity series UHPLC system (Agilent Technologies, Waldbronn, Germany) was used for HILIC-UHPLC experiments with the following conditions: the column Acquity UPLC BEH HILIC (150 × 2.1 mm; 1.7 μm), the column temperature 40°C, the flow rate 0.25 mL/min, and the injection volume 1 μL. The mobile phase A was acetonitrile/water (96/4, v/v), the mobile phase B was acetonitrile/water (50/50, v/v) and both phases contained 15 mM of ammonium acetate. The following gradient elution was used: 0 min, 100 % A; 7.5 min, 84 % A; 11 min, 70 % A; 12 min, 100 % A; 22 min, 100 % A. The total run time including equilibration is 22 min. The autosampler temperature was set at 4°C. The UHPLC system was coupled to a Xevo G2-XS-QTOF mass spectrometer (Waters, Milford, MA, USA). The data were acquired in the sensitivity mode using ESI in the positive and negative ion mode using the following conditions: the capillary voltage of 3 kV for the positive ion mode and 1.5 kV for the negative ion mode, the sampling cone of 20 V, the source offset of 90 V, the source temperature of 150°C, the desolvation temperature of 500°C, the cone gas of 50 L/h, and the desolvation gas flow of 1000 L/h for both ion modes. Mass spectra were measured in the *m/z* range of 150-1500 with the scan time of 0.5 s using the continuum mode and the lock mass scanning. The peptide leucine enkephalin was used as the lock mass for all MS experiments. MS/MS experiments were performed with a collision energy of 35 eV.

The measured data were submitted for noise reduction with the threshold of 20 by the Waters Compression Tool and the lock mass correction was subsequently applied to obtain accurate masses, and the data were converted to the centroid mode. The text files including summarized *m/z* with the corresponding intensities were exported by MarkerLynx XS software. The MS data were processed by a laboratory-made Excel macro script named LipidQuant with an embedded database (70). Lipid species were detected according to accurate *m/z* values with the mass tolerance of 10 mDa. The quantitative analysis was performed using internal standards, and isotopic correction II type was applied. The concentration is expressed in pmol/mg of cells.

The lipids with average concentration lower than 0.1 pmol/mg were excluded for statistical evaluation. Absolute concentrations of lipid species were used for MDA by SIMCA software. The Pareto scaling and logarithm transformation were performed for the data normalisation. Differences were investigated using statistical projection methods PCA, OPLS-DA, and S-plot (**Fig. S2A** and **C**). The quantification data were transformed using a base 2 logarithm. For fold change, the average of the log2 Cas9 control replicates was subtracted from the average of the log2 of the knockout replicates and plotted with standard errors (**Fig. S2B** and **D**).

### Plasma Membrane Profiling (PMP) and Data Processing

#### PMP labelling and streptavidin enrichment

For PMP analysis, 6 x 10^5^ MO3.13 cells were seeded into 15 cm dishes and incubated for 24 hrs in DMEM containing 10% (v/v) FCS. Medium was then replaced with differentiation medium (as described above) and incubated for a further 7 days. PMP was performed as described previously with minor modifications (71, 72). Cells were washed twice in ice-cold PBS. Surface sialic acid residues were oxidised and biotinylated by incubation in the dark for 30 min with gentle rocking at 4°C using an oxidation/biotinylation mix comprising 1 mM sodium meta-periodate, 100 μM aminooxy-biotin (Biotium Inc., Hayward, CA) and 10 μM aniline (Sigma-Aldrich) in ice-cold PBS, pH 6.7. The reaction was quenched with 1 mM glycerol and cells were washed twice in ice-cold PBS. Biotinylated cells were scraped into lysis buffer (10 mM Tris pH 7.6, 1% Triton X-100, 150 mM NaCl, protease inhibitor (complete, without EDTA (Roche), ½ tablet per 20 mL), and 5 mM EDTA) and incubated on a rotating wheel at 4°C for 30 mins. Lysate was cleared by centrifugation (20,000 *g*, 10 mins) at 4°C and supernatant snap frozen in liquid nitrogen and stored at −80°C until further processing. Protein concentrations of thawed samples were determined by bicinchoninic acid (BCA) assay and equal quantities of samples used in subsequent steps. Biotinylated glycoproteins were enriched by incubation for 3 hrs at 4°C on a rotating wheel with high affinity streptavidin agarose beads (ThermoFisher Scientific). Extensive washing was performed on a vacuum manifold, using lysis buffer, then PBS containing 0.5% (w/v) SDS then washed with urea buffer (6 M urea, 50 mM Triethylammonium bicarbonate, TEAB). Beads were incubated at room temperature with shaking for 30 mins in reduction/alkylation mix (urea buffer containing 20 mM iodoacetamide and 10 mM TCEP). Beads were further washed with urea buffer and then 50 mM TEAB and transferred to a fresh tube. Beads were pelleted gently (550 *g*, 2 mins), supernatant removed. Captured protein was digested on-beads in 100 μL of 50 mM TEAB with 1 μg trypsin (Pierce, MS-grade) at 37°C with shaking for 4 hrs. Beads were pelleted gently and supernatant containing trypsinised peptides were dried using a speedvac (Thermo Scientific).

#### TMT Labelling and clean-up

Samples were resuspended in 21 μL 100mM TEAB pH 8.5. After allowing to come to room temperature, 0.5 μg TMTpro (GALC-KO) or 0.2 μg TMT reagents (UGT8-KO) (Thermo Fisher) were resuspended in 9 μL anhydrous acetonitrile (ACN) which was added to the respective samples and incubated at RT for 1 hr. For the GALC-KO experiment, TMTpro reagents were used in ascending order of reporter molecular mass to label: 3× WT, 3× Cas9 only, 3× GALC-KO G1, 3× GALC-KO G2 and 3× GALC rescue. For the UGT8-KO experiment, TMT reagents were similarly used to label: 3× Cas9 only, 3× UGT8-KO G1, 3×UGT8-KO G2. A 3 μL aliquot of each sample was taken and pooled to check TMT labelling efficiency and equality of loading by LC-MS. Samples were stored at −80°C in the interim. After checking each sample was at least 98% TMT labelled, total reporter ion intensities were used to normalise the pooling of the remaining samples such that the final pool should be as close to a 1:1 ratio of total peptide content between samples as possible. This final pool was then dried in a vacuum centrifuge. The sample was acidified to a final 0.1% Trifluoracetic Acid (TFA) (~200μL volume) and Formic Acid (FA) was added until the sodium deoxycholate (SDC) visibly precipitated. Four volumes of ethyl acetate were added and the sample vortexed vigorously for 30s. Sample was then centrifuged at 15,000 *g* for 5 mins at RT to effect phase separation. A gel loading pipette tip was used to withdraw the lower (aqueous) phase to a fresh low adhesion microfuge tube. If any obvious SDC contamination remained the two-phase extraction with ethyl acetate was repeated. The sample was then partially dried in a vacuum centrifuge and brought up to a final volume of 1mL with 0.1% TFA. FA was added until the pH was <2, confirmed by spotting onto pH paper. The sample was then cleaned up by solid phase extraction using a 50 mg tC18 SepPak cartridge (Waters) and a positive pressure manifold (Tecan Resolvex M10). The cartridge was wetted with 1 mL 100% methanol followed by 1 mL ACN, equilibrated with 1 mL 0.1% TFA and the sample loaded slowly. The cartridge was washed with 1 mL 0.1% TFA before eluting with 750 μL 80% ACN, 0.1% TFA. Eluate was dried in a vacuum centrifuge.

#### Basic pH reversed phase fractionation

Samples were resuspended in 40 μL 200 mM ammonium formate pH 10 and transferred to a glass HPLC vial. High pH reverse phase fractionation was conducted on an Ultimate 3000 UHPLC system (Thermo Scientific) equipped with a 2.1 mm × 15 cm, 1.7 μm Kinetex EVO column (Phenomenex). Solvent A was 3% ACN, Solvent B was 100% ACN, solvent C was 200 mM ammonium formate pH 10. Throughout the analysis solvent C was kept at a constant 10%. The flow rate was 500 μL/min and UV was monitored at 280 nm. Samples were loaded in 90% A for 10 min before a gradient elution of 0-10% B over 10 min (curve 3), 10-34% B over 21 min (curve 5), 34-50% B over 5 min (curve 5) followed by a 10 min wash with 90% B. Fractions (100 μL) were collected throughout the run. Fractions containing peptide (as determined by A280) were recombined across the gradient to preserve orthogonality with on-line low pH RP separation. For example, fractions 1, 25, 49, 73, 97 are combined and dried in a vacuum centrifuge and stored at −20°C until LC-MS analysis. Twelve fractions were generated in this way.

#### Mass Spectrometry

Samples were analysed on an Orbitrap Fusion instrument on-line with an Ultimate 3000 RSLC nano UHPLC system (Thermo Fisher). Samples were resuspended in 10 μL 5% DMSO/1% TFA and loaded via a trapping column. Trapping solvent was 0.1% TFA, analytical solvent A was 0.1% FA, solvent B was ACN with 0.1% FA. Samples were loaded onto a trapping column (300 μm × 5mm PepMap cartridge trap; Thermo Fisher) at 10 μL/min for 5 minutes. Samples were then separated on a 75 cm × 75 μm i.d. 2 μm particle size PepMap C18 column (Thermo Fisher). The gradient was 3-10% B over 10 mins, 10-35% B over 155 minutes, 35-45% B over 9 minutes followed by a wash at 95% B for 5 mins and re-equilibration at 3% B. Eluted peptides were introduced by electrospray to the MS by applying 2.1 kV to a stainless steel emitter (5 cm x 30 μm; PepSep). During the gradient elution, mass spectra were acquired with the parameters detailed in **Fig. S14A** using Tune v3.3 and Xcalibur v4.3 (Thermo Fisher). For TMTpro/TMT experiments the isobaric loss exclusion settings were adjusted appropriately.

#### Data Processing

Data were processed with PeaksX+, v10.5 (Bioinfor). Processing parameters are shown in detail in **Fig. S14B**. Briefly, .raw files were searched iteratively in three rounds, with unmatched DeNovo spectra, at 0.1% peptide spectrum match (PSM) false discovery rate (FDR), from the previous search used as the input for the next. The three iterations were as follows: 1) Swissprot Human (27/03/2020) + common contaminants; 2) The same databases as search 1 but permitting semi-specific cleavage; 3) trEMBL Human (27/03/2020), with specific cleavage rules. Proteins were then quantified using the parameters outlined in **Fig. S14C**. Identified proteins and their TMT reporter abundances were output to .csv, imported to R and submitted to statistical analysis using LIMMA, a moderated t-test available through the Bioconductor package (73). LIMMA *p*-values were corrected for multiple hypothesis testing using the Benjamini-Hochberg method to generate an FDR (*q*-value) for each comparison. The mass spectrometry proteomics data have been deposited to the ProteomeXchange Consortium via the PRIDE (74) partner repository with the dataset identifier PXD036727 and 10.6019/PXD036727.

### Plasmids and constructs for protein expression

Amino acid numbering used throughout is based on the following sequences: human NFASC; UniProt ID: 094856, isoform 9 for NF155 and isoform 8 for NF186. Details of the amino acid composition of these isoforms and of the truncation constructs used in this study are provided in **Fig. S5** and **Table S6**.

### Protein expression and purification

Codon-optimised NF155 and NF186 synthetic genes were purchased from GeneArt, and the relevant truncations cloned into the pHLSec expression vector using the *AgeI* and *KpnI* restriction sites. Constructs were sequence verified and then transiently transfected in HEK293-F cells, using the PEI MAX^®^ (Polysciences, #24765) transfection reagent, as per the manufacturer’s instructions. The proteins were expressed for 4 days at 37°C in a humidified 8% CO_2_ atmosphere. The media was harvested by centrifugation at room temperature, beginning with 100 *g* for 5 min to gently pellet and remove the cells, followed by a 4000 *g* for 10 mins to further clarify the media before filtering through a 0.2 μm nylon membrane (Millipore). The media was then incubated with 1.5 mL of Ni-NTA beads (Qiagen) at 4°C on a rolling bed for 1 hr. The Ni-NTA beads were captured using a gravity flow Econo-column (Biorad) and washed with wash buffer containing 50 mM HEPES pH 7.4, 150 mM NaCl and 20 mM imidazole, and protein was eluted in elution buffer containing 50 mM HEPES pH 7.4, 150 mM NaCl and 300 mM imidazole. The eluted protein was further purified via size exclusion chromatography in buffer containing 50 mM HEPES pH 7.4 and 150 mM NaCl. NF186 full-length ECD, NF155 full-length ECD and NF155 IgG1-Fn2 were purified using a HiLoad 16/600 Superdex 200 pg column (Cytiva) and NF155 IgG1-4, NF186 IgG1-4, NF155 IgG5-Fn2 and NF155 Fn1-4 were purified using a HiLoad 16/600 Superdex 75 pg column (Cytiva).

To make deglycosylated full-length NF155-ECD (NF155-DG), HEK293-F cells were transiently transfected in the presence of 5 μM kifunensine and purified as above, to yield the high mannose form (NF155-HM). This was then treated with 1 U Endo Hf (New England Biolabs) per 1 μg of NF155, and incubated at 30°C for 3 hrs. The NF155-DG was separated from the Endo-Hf by loading onto a 5 mL HisTrap™ FF (Cytiva, #17528601) column in wash buffer and then eluting the NF155-DG in elution buffer. NF155-DG was then concentrated to 1.3 mg/mL and buffer exchanged into 50 mM HEPES pH 7.4 and 150 mM NaCl using an Amicon Ultra 30 kDa centrifugal concentrator (Millipore).

### Liposome binding assays

Phosphotidylcholine (PC) (#840051C), sulfatide (#131305P), 1,2-dimyristoyl-sn-glycero-3-phosphoethanolamine-*N*-(lissamine rhodamine B sulfonyl) (Rhod-PE) (#810157P), L-α-phosphatidylserine (PS) (#840032P), D-galactosyl-α-1,1’ N-palmitoyl-D-erythro-sphingosine (GalCer) were purchased from Avanti. Ganglioside GM_4_ (GM_4_) was purchased from Merck (#345748). Lipids were dissolved/suspended in chloroform (aside from GalCer, which was dissolved in 2:1 chloroform:methanol) and the final desired mixtures added to a glass vial before the solvent was evaporated under a nitrogen stream. To all liposomes prepared, 2% Rhod-PE was incorporated to aid with liposome visualisation. The resultant lipid film was then hydrated using liposome buffer (50 mM HEPES pH 7.4 and 150 mM NaCl) to form multilamellar liposomes. A final liposome concentration of 0.8 mM in 50 μL was incubated with 1 μM protein for 30 min at room temperature with rotation. The liposomes were then pelleted by centrifugation at 20,000 *g* and the supernatant removed. A further 50 μL aliquot of liposome buffer was added to the pellet and vortexed to wash the sample before a second centrifugation. The supernatant was removed once more and the pellet resuspended in 10 μL of liposome buffer and 10 μL of 2× loading dye, separated using a NuPAGE 4-12% Bis-Tris gel (Invitrogen), stained with InstantBlue (Abcam) and imaged using a BioRad ChemiDoc MP imaging system. Densitometric analysis of gel images was performed using ImageJ and GelAnalyzer. For fitting of binding data, n=4 independent experiments comparing binding of IgG1-Fn4 and IgG1-Fn2 were performed. For each replicate the percentage of bound NF155 was fitted to the Hill equation using Prism version 9 (GraphPad), yielding values for B_max_, the Hill coefficient and the percentage of sulfatide for half-maximal binding. Statistical tests to compare values for IgG1-Fn4 and IgG1-Fn2 were performed using a two-tailed paired t-test in Prism, to control for liposome variability between experiments.

### Multi-angle Light Scattering (MALS)

MALS experiments were performed immediately following SEC (SEC-MALS) by inline measurement of static light scattering (DAWN 8+; Wyatt Technology), differential refractive index (Optilab T-rEX; Wyatt Technology), and UV absorbance (1260 UV; Agilent Technologies). 1 mg/mL samples of NF155 constructs (100 μL) were injected onto an Superdex 200 Increase 10/300 GL column (Cytiva) equilibrated in in 50 mM HEPES pH 7.4, 150 mM NaCl at 0.4 mL/min. The molar masses of the major SEC elution peaks were calculated in ASTRA 6 (Wyatt Technology) using a protein dn/dc value of 0.185 mL/g. For determination of protein and glycan fractions, conjugate analysis was performed in ASTRA 6, using a glycan (modifier) dn/dc = 0.14 mL/g and theoretical UV extinction coefficient calculated using ProtParam (75). Figures were prepared using Prism 9 (GraphPad).

### Negative stain electron microscopy

Purified NF155-ECD (10 nM) in 50 mM HEPES pH 7.4 and 150 mM NaCl was applied to glow-discharged 300 mesh formvar/carbon coated copper grids (Agar Scientific) and stained with 2% (w/v) uranyl acetate using the side blotting method, as previously described (76). Electron micrograph images were collected using a FEI Tecnai G2 Spirit BioTWIN transmission electron microscope operating at 120 kV, equipped with an Ultrascan 1000X-U CCD camera (Gatan). Data collection was performed at 30,000 × nominal magnification (pixel size 3.31 Å) with a total electron dose of 20-40 e^-^/ Å^2^ and −1.5 μm nominal defocus across 1 s exposures. In total, 95 images were taken and analysed using CryoSPARC version 3.2.0. Images were CTF corrected, and 1000 particles were picked manually, followed by template picking with an extraction box size of 150 pixels. From this, 26,642 particles were picked, generating 28 2D classes that were used for *ab initio* reconstruction of the NF155-ECD. This was followed by homogenous refinement with an initial low pass resolution of 50 Å and a maximum align resolution of 10 Å. The overall resolution of the generated 3D map was calculated using the CryoSPARC Local Resolution tool with a Fourier Shell Correlation (FSC) cut-off of 0.143 (77).

### Generating AF2 models and fitting to the EM map

Structural models were generated using a locally installed version of ColabFold version 1.3, implementing AlphaFold2 machine learning structure prediction, with default parameters (50, 51). AF2 models for the FL-ECD were positioned by rigid-body docking into the three-dimensional EM density map using the ‘fit in map’ tool in UCSF ChimeraX (78). The C-terminal tail of the FL-ECD was removed from the model because it was predicted to be disordered (pLDDT < 50) and would thus not be resolved in the EM map. Correlation coefficients were calculated by generating maps around the docked atomic models, at the resolution of the EM map, and calculating map-to-map correlations in ChimeraX. Improved fitting of models to the EM map was carried out using molecular dynamics with flexible fitting (MDFF) as implemented by ISOLDE within ChimeraX (53). MDFF simulations used the AF2 model as a reference structure with strong distance and torsion restraints weighted using the PAE matrix and pLDDT values of the AF2 model, respectively. Following visual inspection of the model docked into the EM map, the IgG1-4 domain was manually repositioned by rotating to maximise the fit to density and MDFF simulations were repeated as above, maintaining the AF2 model as reference and retaining strong distance and torsion restraints. All models and associated statistics will be deposited in the University of Cambridge Data Repository (TBA).

### Fluorescent liposome clustering assay

Liposomes were prepared as detailed above. Additional lipids, 1,2-dioleoyl-sn-glycero-3-[(N-(5-amino-1-carboxypentyl)iminodiacetic acid)succinyl] (NiNTA-DGS) (#790404P) and 1,2-dimyristoyl-sn-glycero-3-phosphoethanolamine-N-(7-nitro-2-1,3-benzoxadiazol-4-yl) (NBD-PE) (#810143P), were purchased from Avanti^®^ and dissolved in chloroform. Three lipid compositions (molar percentage) were made: 1) 4% NiNTA-DGS, 2% NBD and 94% PC, 2) 50% sulfatide, 4% NiNTA-DGS, 2% NBD-PE and 44% PC, 3) 50% sulfatide, 2% Rhod-PE, 48%. Samples containing 0.4 mM liposomes with compositions 1 or 2 (as detailed above) were preincubated with 20 nM NF155-ECD in liposome buffer at room temperature with rotation for 15 min. Liposome composition 3 was added (0.4 mM) to each and incubated for a further 15 min. Samples were diluted 10-fold and placed on a microscope slide, under a cover slip. Liposomes were imaged using the EVOS M5000 Imaging System at 20 x magnification. Colocalisation of the pink and green liposomes was quantified using the ImageJ coloc2 software.

## Acknowledgements

This work was supported by a Royal Society University Research Fellowship (UF100371) to JED and a Wellcome Trust Senior Research Fellowship (219447/Z/19/Z) to JED that also supports SJM, ASN and ES. ERC was supported by a Royal Society Enhancement Award to JED (RGF\EA\180151) and SF was supported by an MRC Project Grant to JED (MR/N020626/1). JCW was supported by a Wellcome Trust Principal Research Fellowship to PJL (210688/Z/18/Z). BGB was supported by a Wellcome Trust PhD studentship from the University of Cambridge School of Clinical Medicine. MH acknowledges the support of grant project No. 21-20238S sponsored by the Czech Science Foundation.

## Data Availability

The mass spectrometry proteomics data produced in this study have been deposited to the ProteomeXchange Consortium via the PRIDE repository (74) (https://www.ebi.ac.uk/pride/) with the dataset identifier PXD036727 and 10.6019/PXD036727. Lipidomics data are supplied with this submission as a supplementary file. AF2 models and associated statistics will be deposited in the University of Cambridge Data Repository (TBA). Raw electron microscopy images, processed data and final models will be deposited in the University of Cambridge Data Repository (TBA) and the Electron Microscopy Public Image Archive EMPIAR (79).

## Notes

### Competing Interest Statement

The authors have declared no competing interest.

### Summary of Updates

Additional data in 2D to support change in affinity but not Hill co-efficient following removal of C-terminal Fn domains of NF155.

