## Supplementary Figures and Tables for "Altered plasma membrane abundance of the sulfatide-binding protein NF155 links glycosphingolipid imbalances to demyelination"

### Supplementary Data

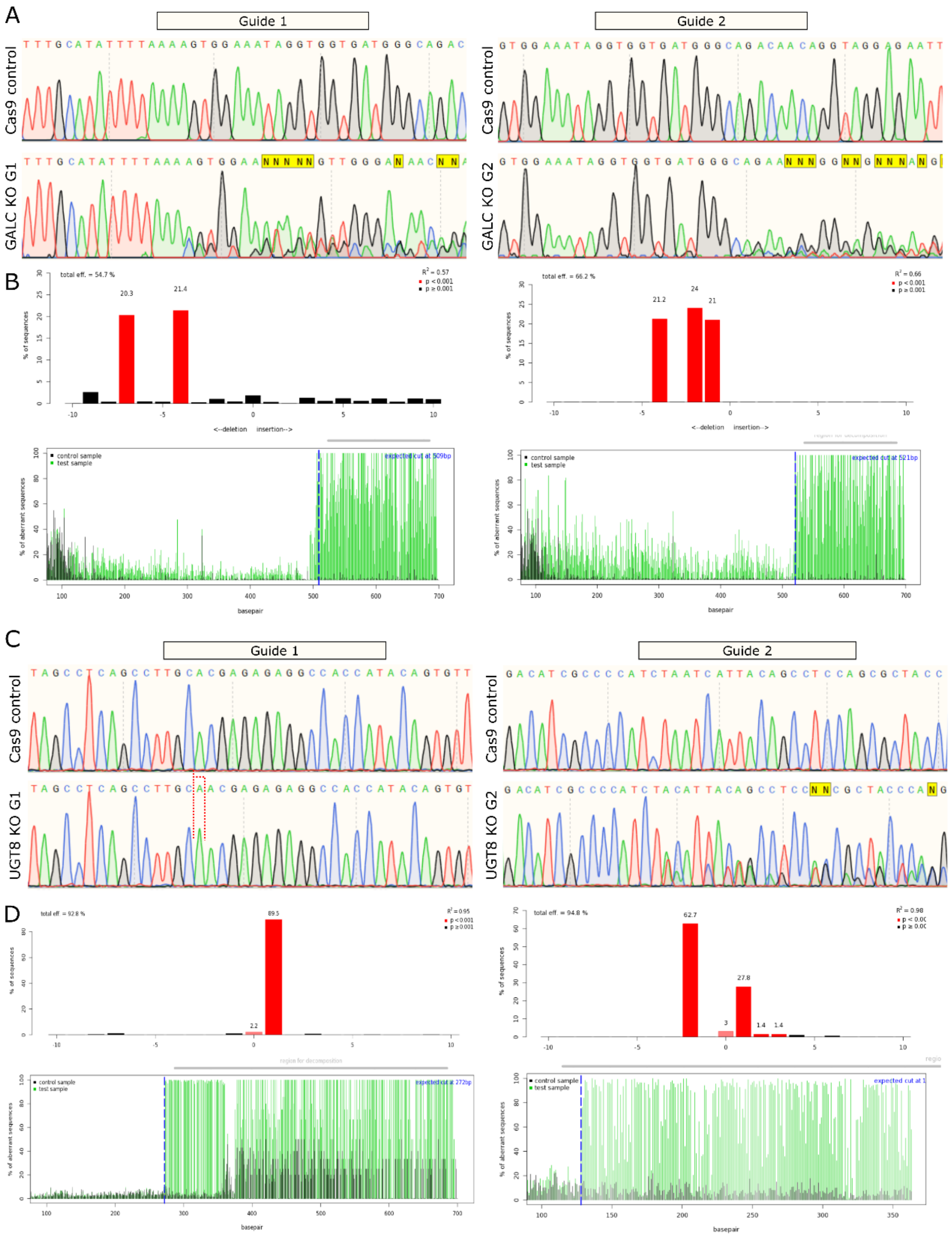


**Figure S1. Confirmation of editing at gRNA-targeted sites of GALC and UGT8 genes. (A)** DNA sequencing chromatograms are shown across the relevant region of the genome for cell lines transfected with untargeted Cas9 (Cas9 control, *top*), showing the unedited sequence, and for Cas9 targeted by sgRNA1 and sgRNA2 to the GALC gene (*bottom*) in the clonal MO3.13 lines used in this study. Overlapping, altered sequences demonstrate editing at the target site. **(B)** Analysis of sequence chromatograms using TIDE (1), predicting the frequency and position of deletions and insertions as a result of CRISPR/Cas9 editing. **(C and D)** As for panels (A and B) but for the UGT8 KO clonal cell lines.

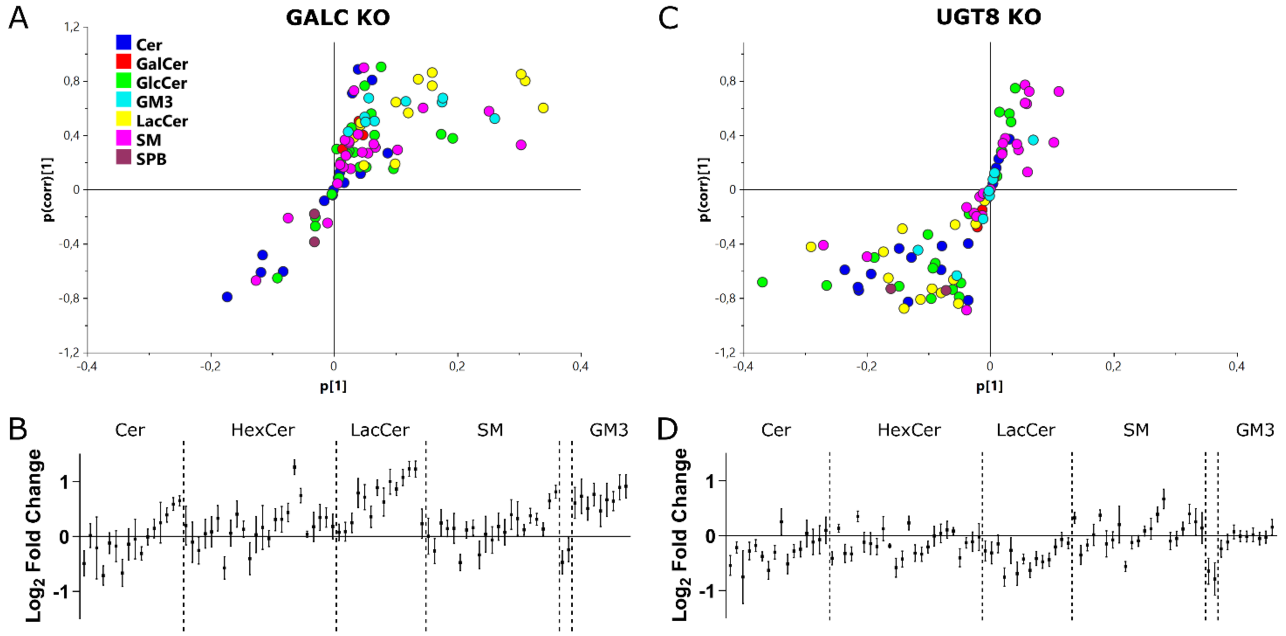


**Figure S2. Targeted lipidomics quantifying ceramide, sphingomyelin and glycosylated sphingolipids. (A)** S-plot demonstrating the up-regulated (upper right corner) and down-regulated (lower left corner) sphingolipid species comparing GALC KO samples (n=3 for each of two cell lines) with Cas9 control cells (n=3). **(B)** The same lipidomics data as in (A) analysed by lipid class. The log base-2 fold change in abundance between GALC KO cells and Cas9 control cells of all quantified sphingolipids. Samples (n=3) for each of the two GALC KOs were included in the analysis against Cas9 cells (n=3). Mean and standard error are plotted for each lipid species. **(C)** S-plot analysis as in (A) for UGT8 KO cell lines. **(D)** Quantification by lipid species as in (B) for UGT8 KO cell lines.

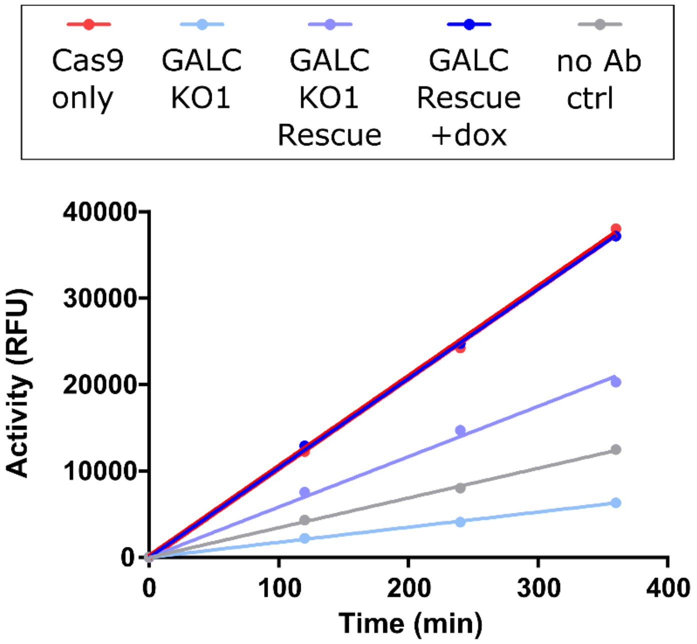


**Figure S3. Activity assays for the control, GALC KO and GALC rescue cell lines.** GALC activity assays following immunoprecipitation (IP) from control, GALC KO1 and GALC KO1 Rescue cell lines in the absence and presence of doxycycline (GALC expression in the rescue cell line is controlled by a doxycycline-inducible promoter). Low levels of GALC activity are present in the rescue cell line in the absence of dox, but with the addition of 0.2 µg/mL dox GALC expression is restored to control (Cas9 only) activity levels. A no antibody (no Ab) control IP is also included.

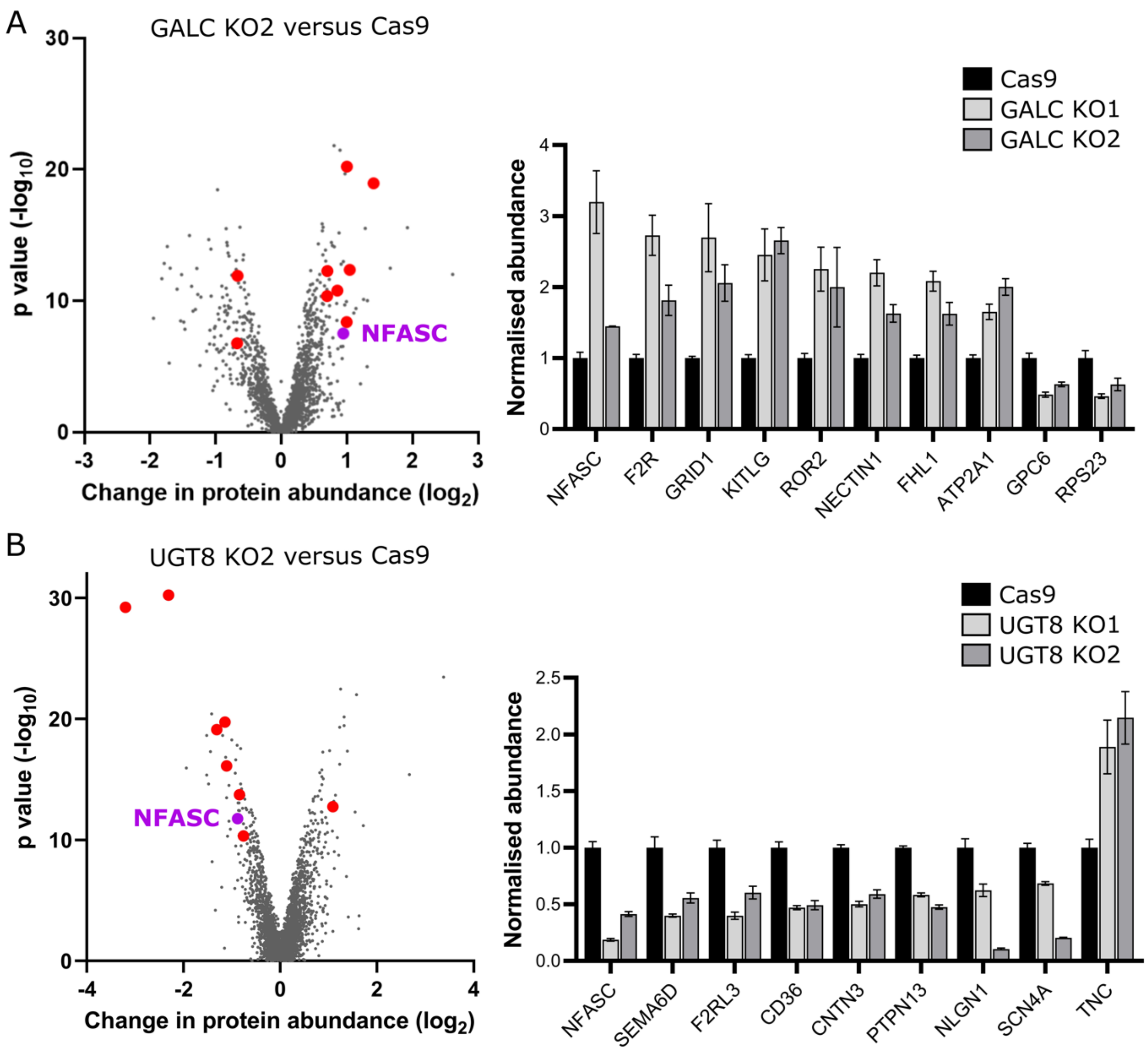


**Figure S4. PMP-MS data for additional clonal GALC and UGT8 KO cell lines. (A)** Quantitative mass spectrometry following enrichment of plasma membrane proteins (PMP-MS) from the GALC KO2 cell line compared with the Cas9 control. *Left*: volcano plot with the horizontal axis showing average fold change across three biological replicates and the vertical axis showing significance (two-sided t test) across the three replicates. The ten high-confidence targets (criteria as detailed in the main text) are coloured in red with NFASC specifically highlighted in purple. *Right*: Normalised protein abundance values, from the PMP-MS data, for high-confidence targets in both the GALC clonal KO cell lines. Data shown in Table S1. Normalised abundances have been used to allow comparison across genes with different absolute abundances in the cell. **(B)** As for panel (A) but for the UGT8 KO2 cell line. Data shown in Table S2.

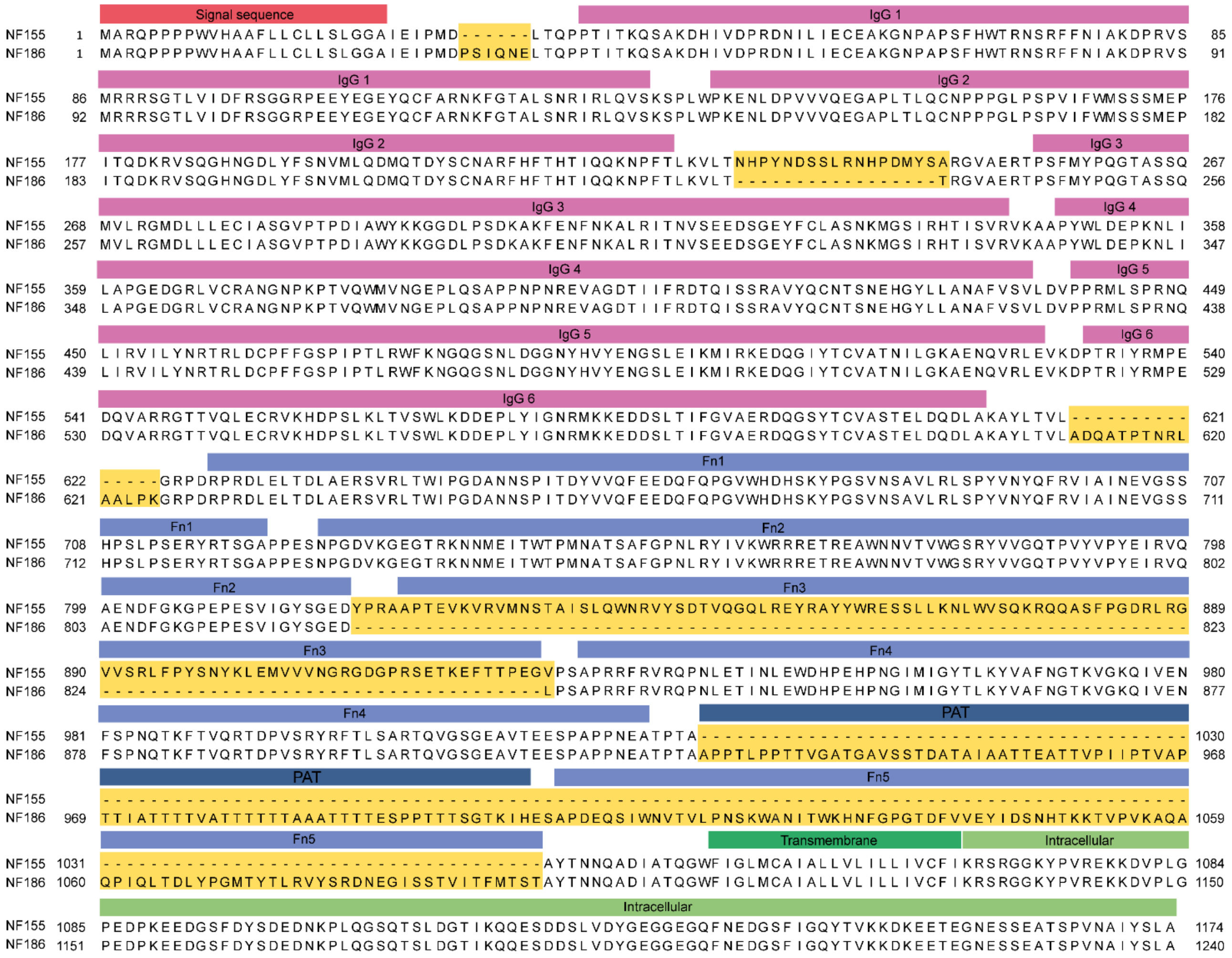


**Fig. S5. Sequence composition of NF155 and NF186 isoforms.** IgG-like domains highlighted in pink, Fn-like domains in light blue, PAT domain in dark blue, transmembrane domain in dark green, intracellular domain in light green and differences between the two isoforms in yellow. Alignment generated using Jalview (2) and annotated in Adobe Illustrator.

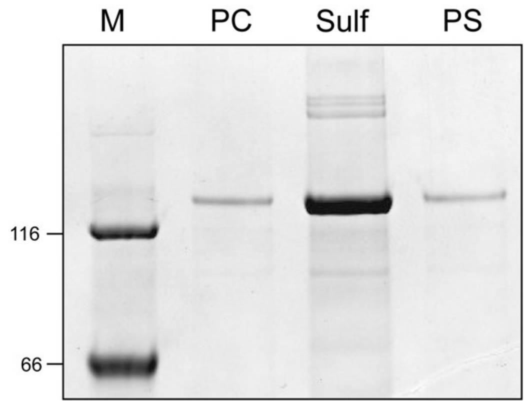


**Fig. S6. NF155 does not bind the anionic lipid phosphatidylserine (PS).** Liposome pull down assay demonstrating that NF155-ECD does not non-specifically bind negatively charged membrane, but specifically binds sulfatide. Here, three liposome compositions were tested for binding to 1 µM NF155 FL ECD: 98% PC: 2% Rhod-PE (lane PC), 28% PC: 2% Rhod-PE: 70% sulfatide (Sulf), and 28% PC: 2% Rhod-PE: 70% PS (lane PS). Samples separated by SDS PAGE and stained using Coomassie.

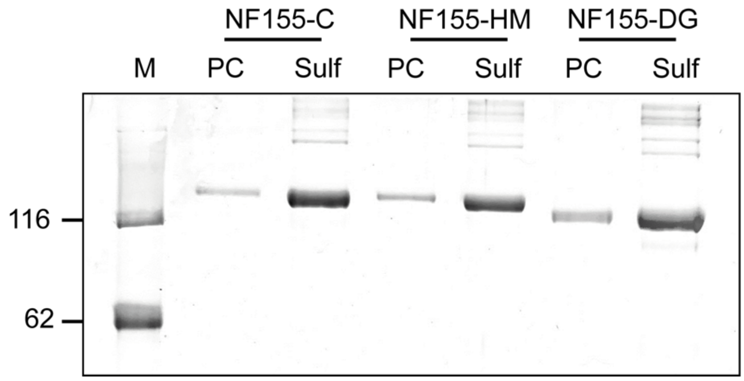


**Fig S7. Deglycosylated NF155 retains binding to sulfatide.** Liposome pull down assay demonstrating that the glycosylation of NF155 FL ECD does not affect sulfatide binding. Liposomes used to test binding of 1 µM NF155 FL ECD were 98% PC: 2% Rhod-PE (PC), or 48% PC: 2% Rhod-PE: 50% sulfatide (Sulf). NF155-C has complex glycosylation, NF155-HM has high mannose glycosylation and NF155-DG is deglycosylated. Samples separated by SDS PAGE and stained using Coomassie. Due to the low stability of NF155 when deglycosylated this experiment was performed immediately following EndoH treatment.

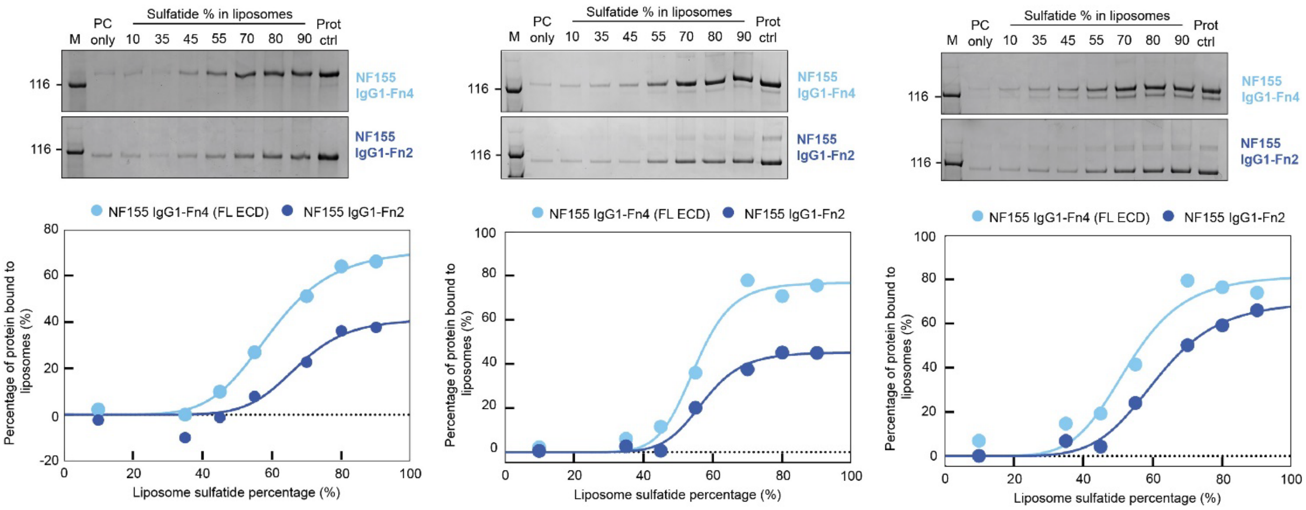


**Figure S8. Replicates of quantitive liposome binding data for NF155 IgG1-Fn4 and IgG1-Fn2.** Three independent replicates of the data shown in Fig. 2D of the main text are displayed as follows. *Top:* Liposome binding assay performed with 250 nM NF155 using 1.6 mM liposomes containing increasing concentrations of sulfatide (IgG1-Fn4, top gel, IgG1-Fn2, lower gel). *Bottom:* Densitometric analysis of the SDS PAGE reveals a sigmoidal binding relationship for both NF155 constructs (IgG1-Fn4, light blue, IgG1-Fn2, dark blue). Prot ctrl represents total protein added to each assay sample. A total of n=4 independent experiments were performed.

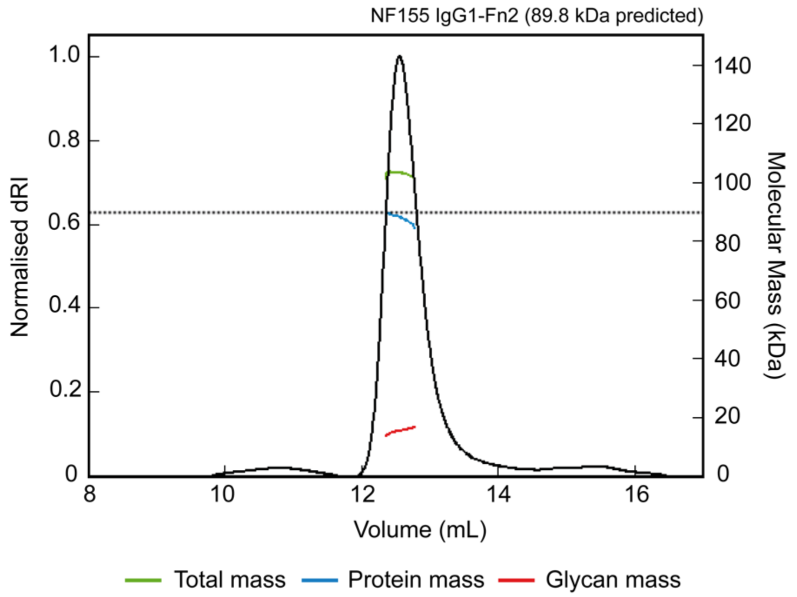


**Fig. S9. Size exclusion chromatography coupled to multi-angle light scattering (SEC-MALS) for the NF155 truncation IgG1-Fn2.** Protein conjugate mass analysis of the elution peak demonstrates that NF155 IgG1-Fn2 is monomeric with a calculated protein mass of 87.6 kDa (consistent with the predicted mass of 89.8 kDa, dotted line) and 15.3 kDa glycans.

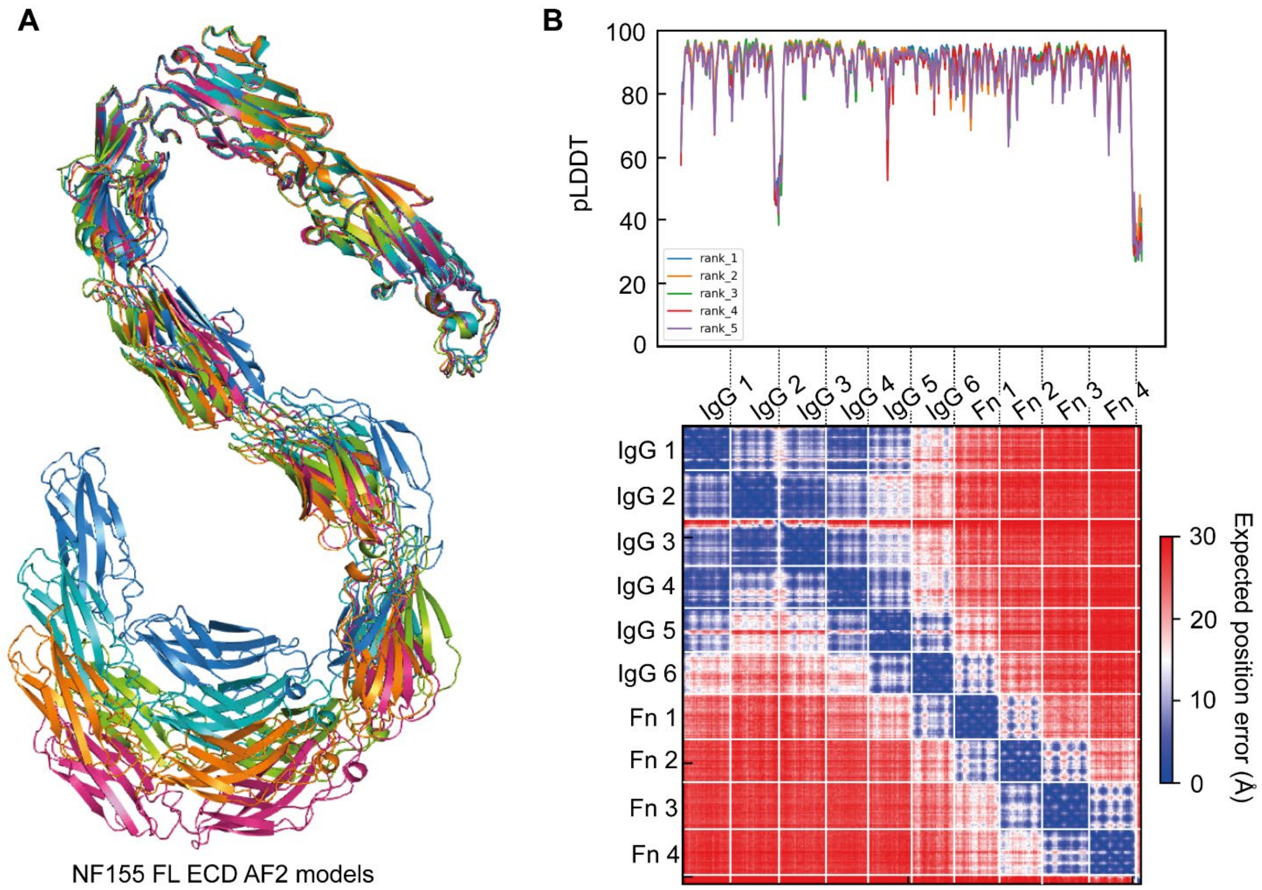


**Fig. S10. AF2 models of the NF155-ECD. (A)** Ribbon diagrams of the five AF2 predictions of the NF155 FL ECD aligned using the four N-terminal domains, IgG1-IgG4. **(B)** *Top,* pLDDT plots demonstrating good per-residue confidence for all five NF155 FL ECD predictions. *Bottom*, Representative Predicted Aligned Error (PAE) plot for NF155 FL ECD, with domain boundaries marked (white lines), demonstrating the low confidence (high error, red) of long-distance predictions, aside from the four N-terminal domains, IgG1-IgG4.

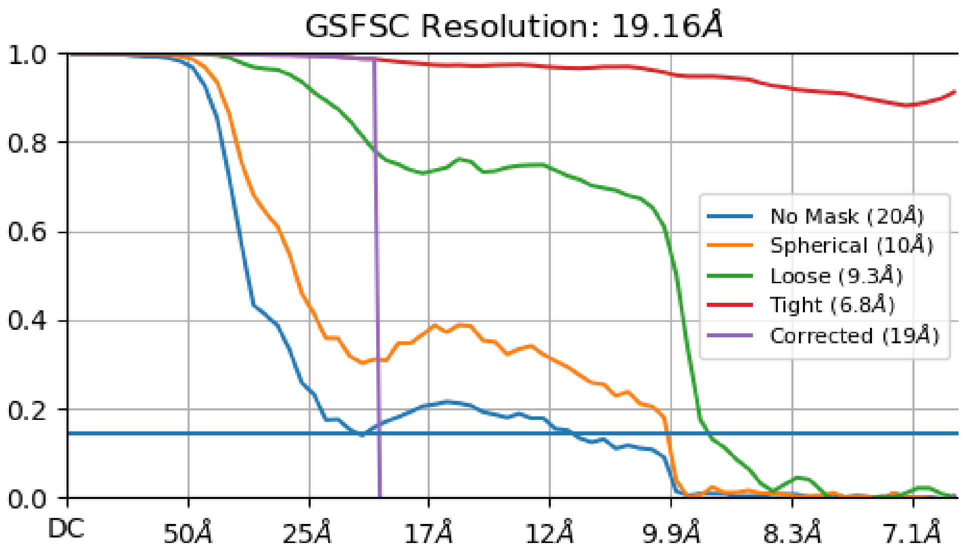


**Fig. S11.** Resolution estimation of the 3D reconstruction of NF155-ECD based on the Fourier Shell Correlation (FSC)=0.143 criterion with different masking options (3). Performed using the CryoSPARC Local Resolution tool.

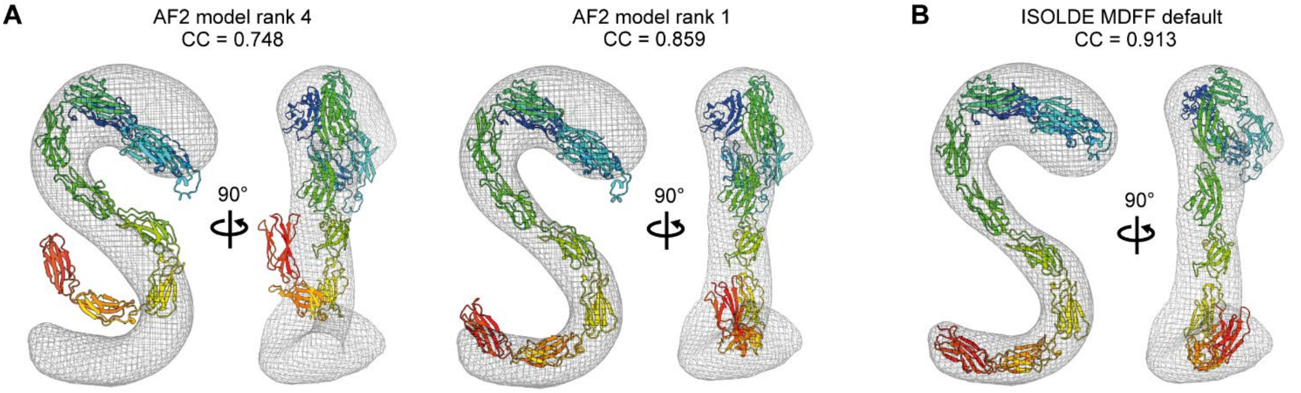


**Fig. S12. Initial docking of AF2 models of the NF155-ECD into negative-stain EM maps. (A)** Rigid-body fits into the negative stain EM map of AF2 models with the lowest (*left*) and highest (*right*) correlation coefficient (CC) when compared to the experimental map. Fit and CC calculations performed using Chimera-X (4). **(B)** The AF2 model rank 1 following molecular dynamics flexible fitting (MDFF) implemented using ISOLDE (5) with strong torsion and distance restrains weighted by the AF2 model pLDDT and PAE scores, respectively. For each representative image two orientations are shown rotated by 90° about the y-axis demonstrating the overall fit to the map.

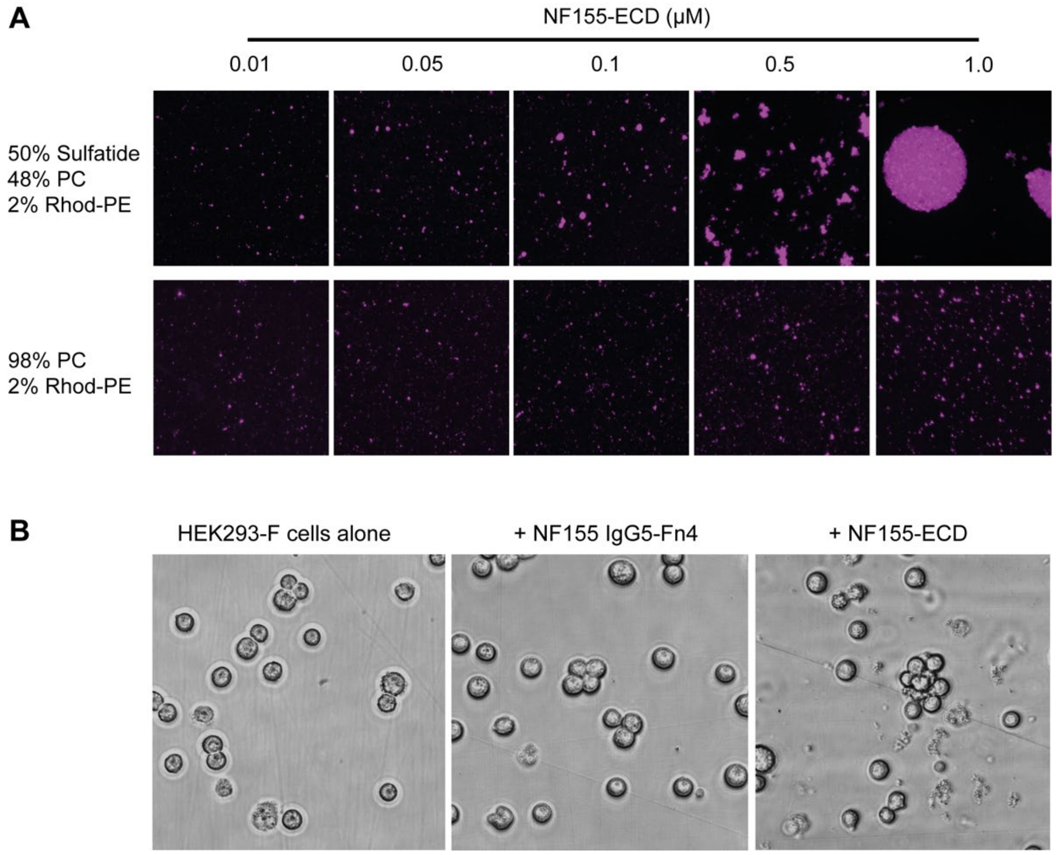


**Fig. S13. NF155-ECD causes liposome aggregation and cellular membrane disruption. (A)** Incubation of increasing concentrations of NF155-ECD with sulfatide-containing liposomes, caused increasingly severe liposome aggregation, while incubation with liposomes without sulfatide had no effect. **(B)** Incubation of NF155-ECD with HEK293-F cells caused increased cell clustering, membrane blebbing and loss of intracellular material, while incubation with the NF155 N-terminal truncation (IgG5-Fn4) did not.

**A** Mass spectrometry method

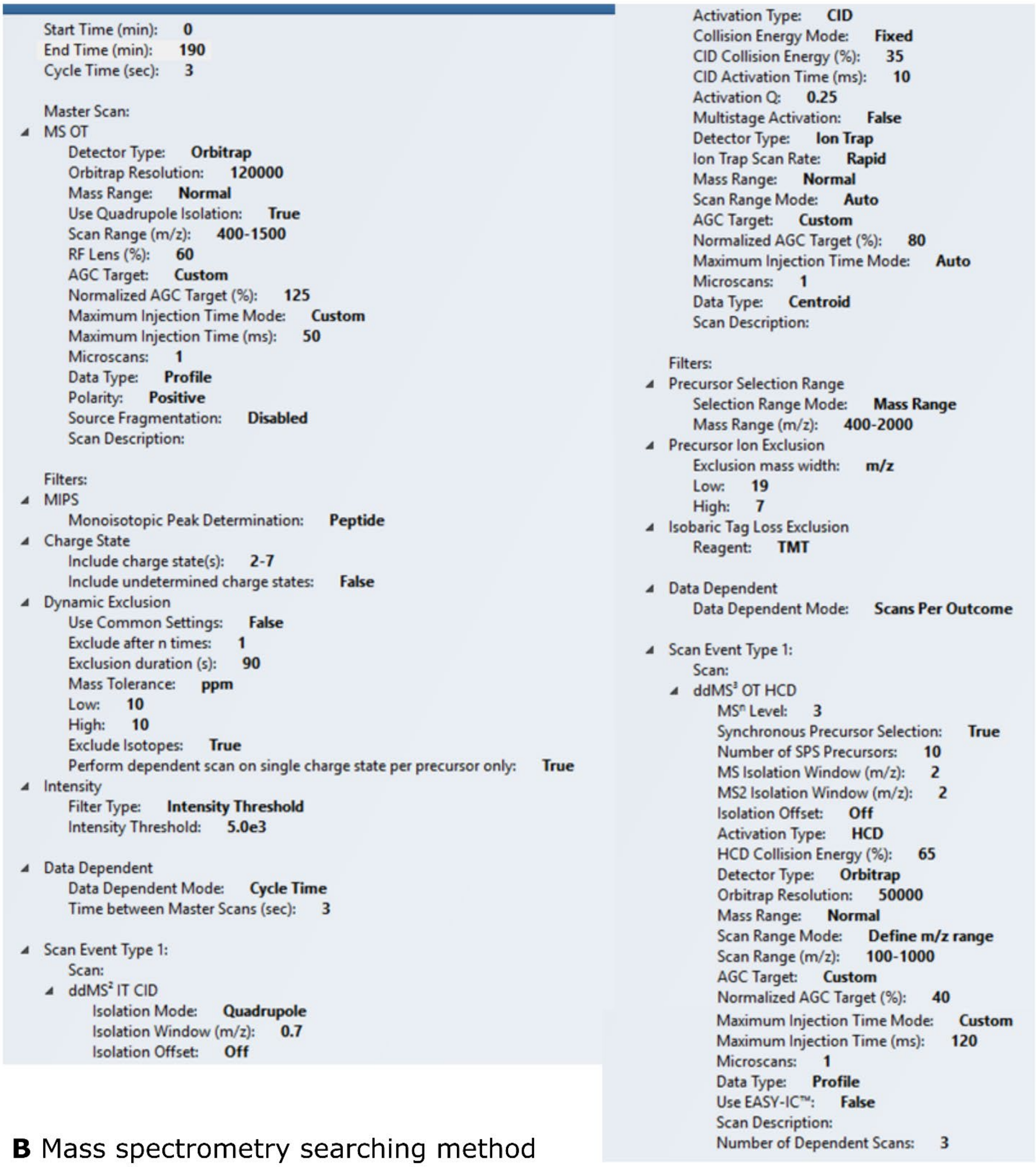

Start Time (min): **0**
End Time (min): **190**
Cycle Time (sec): **3**

Master Scan:
- MS OT
  - Detector Type: **Orbitrap**
  - Orbitrap Resolution: **120000**
  - Mass Range: **Normal**
  - Use Quadrupole Isolation: **True**
  - Scan Range (m/z): **400-1500**
  - RF Lens (%): **60**
  - AGC Target: **Custom**
  - Normalized AGC Target (%): **125**
  - Maximum Injection Time Mode: **Custom**
  - Maximum Injection Time (ms): **50**
  - Microscans: **1**
  - Data Type: **Profile**
  - Polarity: **Positive**
  - Source Fragmentation: **Disabled**
  - Scan Description:

Filters:
- MIPS
  - Monoisotopic Peak Determination: **Peptide**
- Charge State
  - Include charge state(s): **2-7**
  - Include undetermined charge states: **False**
- Dynamic Exclusion
  - Use Common Settings: **False**
  - Exclude after n times: **1**
  - Exclusion duration (s): **90**
  - Mass Tolerance: **ppm**
  - Low: **10**
  - High: **10**
  - Exclude Isotopes: **True**
  - Perform dependent scan on single charge state per precursor only: **True**
- Intensity
  - Filter Type: **Intensity Threshold**
  - Intensity Threshold: **5.0e3**

- Data Dependent
  - Data Dependent Mode: **Cycle Time**
  - Time between Master Scans (sec): **3**

- Scan Event Type 1:
  - Scan:
  - $ddMS^2$ IT CID
    - Isolation Mode: **Quadrupole**
    - Isolation Window (m/z): **0.7**
    - Isolation Offset: **Off**
    - Activation Type: **CID**
    - Collision Energy Mode: **Fixed**
    - CID Collision Energy (%): **35**
    - CID Activation Time (ms): **10**
    - Activation Q: **0.25**
    - Multistage Activation: **False**
    - Detector Type: **Ion Trap**
    - Ion Trap Scan Rate: **Rapid**
    - Mass Range: **Normal**
    - Scan Range Mode: **Auto**
    - AGC Target: **Custom**
    - Normalized AGC Target (%): **80**
    - Maximum Injection Time Mode: **Auto**
    - Microscans: **1**
    - Data Type: **Centroid**
    - Scan Description:

  Filters:
  - Precursor Selection Range
    - Selection Range Mode: **Mass Range**
    - Mass Range (m/z): **400-2000**
  - Precursor Ion Exclusion
    - Exclusion mass width: **m/z**
    - Low: **19**
    - High: **7**
  - Isobaric Tag Loss Exclusion
    - Reagent: **TMT**

  - Data Dependent
    - Data Dependent Mode: **Scans Per Outcome**

  - Scan Event Type 1:
    - Scan:
    - $ddMS^3$ OT HCD
      - $MS^n$ Level: **3**
      - Synchronous Precursor Selection: **True**
      - Number of SPS Precursors: **10**
      - MS Isolation Window (m/z): **2**
      - MS2 Isolation Window (m/z): **2**
      - Isolation Offset: **Off**
      - Activation Type: **HCD**
      - HCD Collision Energy (%): **65**
      - Detector Type: **Orbitrap**
      - Orbitrap Resolution: **50000**
      - Mass Range: **Normal**
      - Scan Range Mode: **Define m/z range**
      - Scan Range (m/z): **100-1000**
      - AGC Target: **Custom**
      - Normalized AGC Target (%): **40**
      - Maximum Injection Time Mode: **Custom**
      - Maximum Injection Time (ms): **120**
      - Microscans: **1**
      - Data Type: **Profile**
      - Use EASY-IC™: **False**
      - Scan Description:
      - Number of Dependent Scans: **3**

**B** Mass spectrometry searching method

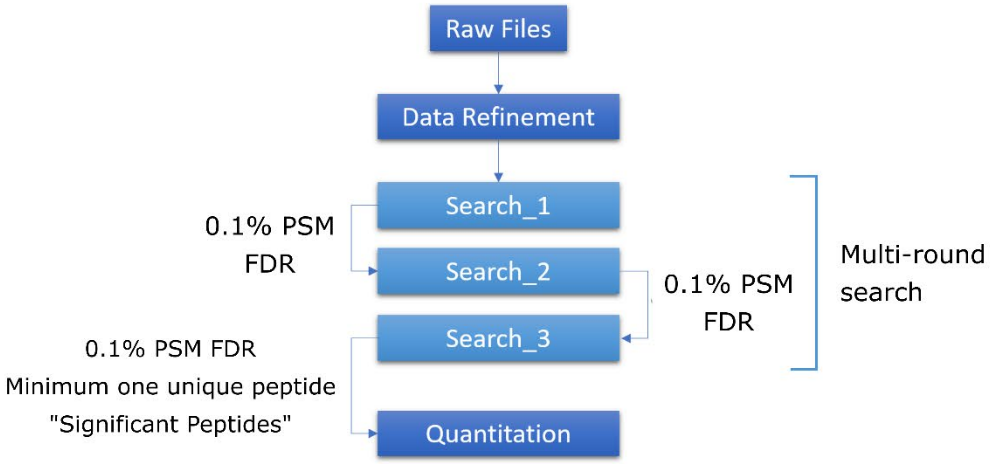

**B** Mass spectrometry searching method (continued)

**Data refinement**

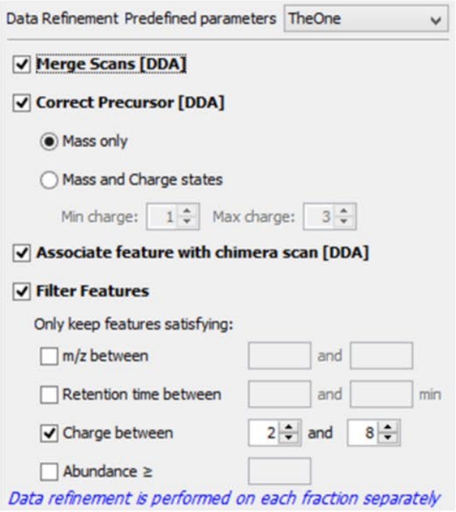


**Cleavage Rules**

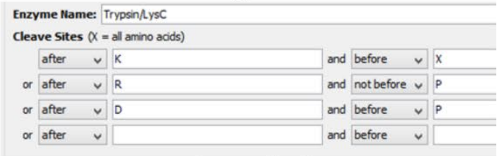


**Search 1**

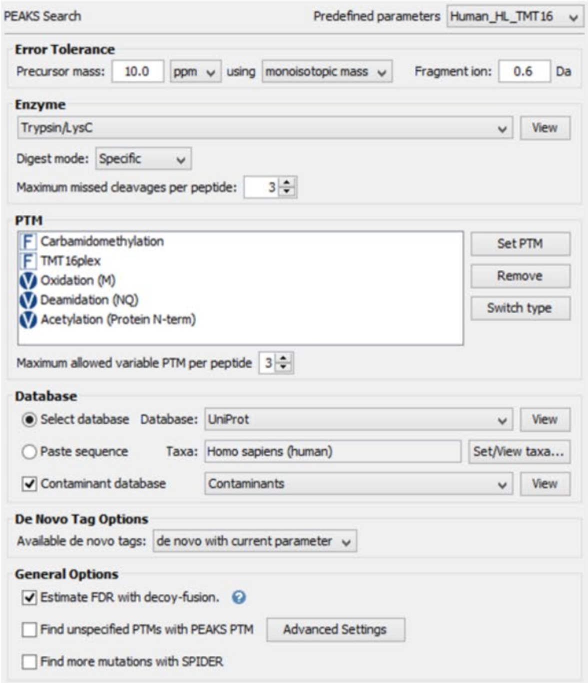


**Search 2**

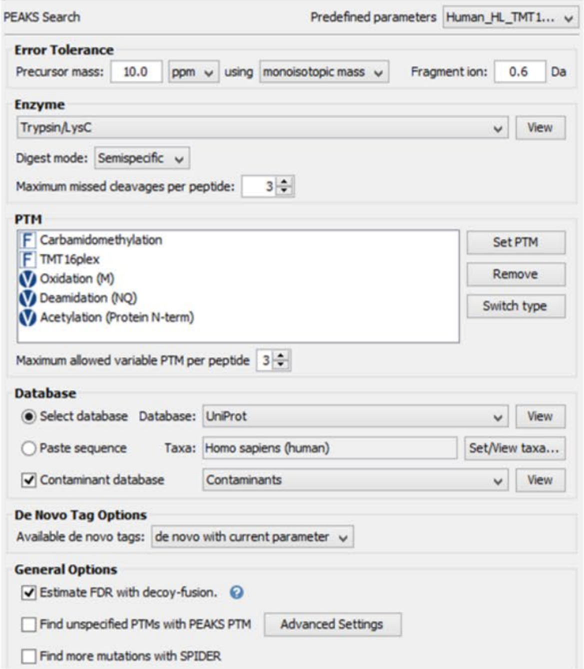


**Search 3**

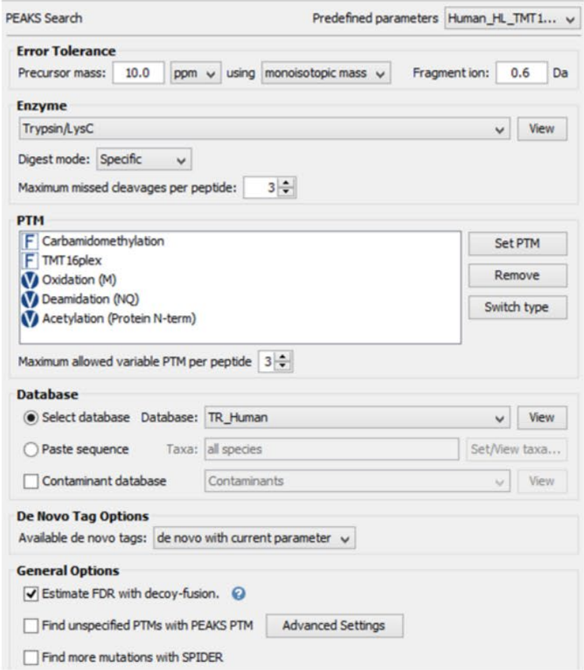


**C** Mass spectrometry quantitation

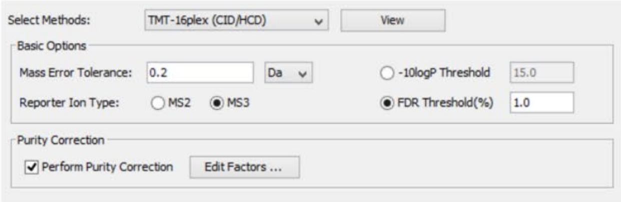


**Fig. S14. Collection and processing of protein mass spectrometry data.** Parameters for protein mass spectrometry data collection **(A)**, data searching **(B)** and quantitation **(C)** for data shown in **Figure 1**.

**Table S1. High confidence targets identified in PMP-MS of GALC cell lines compared with Cas9 control cells.**

| Gene ID | Description | #Unique peptides | GALC KO1/Cas9 | | GALC KO2/Cas9 | | GALC Rescue/GALC KO1 | |
|---|---|---|---|---|---|---|---|---|
| | | | Log2 Fold change | Significance q-value | Log2 Fold change | Significance q-value | Log2 Fold change | Significance q-value |
| NFASC | Neurofascin | 25 | 1.677 | 1.12E-05 | 0.535 | 2.40E-02 | -0.608 | 2.39E-02 |
| F2R | Proteinase-activated receptor 1 | 3 | 1.449 | 1.13E-04 | 0.858 | 6.80E-03 | -1.027 | 2.85E-03 |
| GRID1 | Glutamate receptor delta-1 | 12 | 1.430 | 1.34E-04 | 1.046 | 3.51E-03 | -1.256 | 9.60E-04 |
| KITLG | Kit ligand | 5 | 1.295 | 1.56E-04 | 1.410 | 2.36E-04 | -1.273 | 7.62E-04 |
| ROR2 | Tyr-kinase TM receptor | 6 | 1.171 | 4.83E-03 | 0.999 | 2.02E-02 | -0.907 | 3.61E-02 |
| NECTIN1 | Nectin-1 | 17 | 1.139 | 4.50E-05 | 0.705 | 3.63E-03 | -0.938 | 7.61E-04 |
| FHL1 | Four and a half LIM domain 1 | 2 | 1.059 | 3.91E-04 | 0.699 | 8.39E-03 | -1.073 | 9.88E-04 |
| ATP2A1 | Sarcoplasmic/ER Ca ATPase 1 | 19 | 0.723 | 5.60E-04 | 1.001 | 1.36E-04 | -0.474 | 1.31E-02 |
| GPC6 | Glypican-6 | 4 | -1.045 | 8.75E-05 | -0.665 | 4.18E-03 | 0.503 | 2.41E-02 |
| RPS23 | 40S ribosomal protein S23 | 3 | -1.110 | 2.11E-03 | -0.672 | 4.61E-02 | 0.937 | 1.00E-02 |

**Table S2. High confidence targets identified in PMP-MS of UGT8 KO cell lines compared with Cas9 control cells.**

| Gene ID | Description | #Unique peptides | UGT8 KO2/Cas9 | | UGT8 KO3/Cas9 | |
|---|---|---|---|---|---|---|
| | | | Log2 Fold change | Significance q-value | Log2 Fold change | Significance q-value |
| SEMA6D | Semaphorin-6D | 3 | -1.310 | 4.11E-04 | -2.530 | 1.72E-05 |
| NFASC | Neurofascin | 18 | -0.880 | 1.44E-02 | -1.390 | 1.72E-03 |
| F2RL3 | Proteinase-activated receptor 4 | 3 | -0.760 | 2.57E-02 | -1.350 | 1.70E-03 |
| CD36 | Platelet glycoprotein 4 | 13 | -1.110 | 1.79E-03 | -1.080 | 1.74E-03 |
| CNTN3 | Contactin-3 | 5 | -0.840 | 6.26E-03 | -1.060 | 1.72E-03 |
| PTPN13 | Tyr phosphatase non-receptor 13 | 12 | -1.140 | 3.54E-04 | -0.770 | 2.71E-03 |
| NLGN1 | Neuroligin-1 | 4 | -3.200 | 3.37E-06 | -0.620 | 4.42E-02 |
| SCN4A | Sodium channel type 4 subunit a | 13 | -2.310 | 3.36E-06 | -0.590 | 1.63E-02 |
| TNC | Tenascin | 4 | 1.090 | 9.72E-03 | 0.840 | 2.82E-02 |

**Table S3.** sgRNA target sequences

| Gene | Exon | Target sequence |
|---|---|---|
| **GALC** | 3 | 5' - AAAGTGGAAATAGGTGGTGA |
| **GALC** | 3 | 5' - GGTGGTGATGGGCAGACAAC |
| **UGT8** | 2 | 5' - ATGGTGGCCTCTCTCGTGCA |
| **UGT8** | 2 | 5' - TGGAGGCTGTAATGATTAGA |

**Table S4.** Sequencing primers across genomic edit sites

| Gene | Fwd Primer 5' -> 3' | Rev Primer 5' -> 3' |
|---|---|---|
| **GALC** | GCCCTACTTGCCCAATGATTGGCACAGAAAG | CAAAACTGTCCCTATGCTCTGTCCTGTATATATAG |
| **UGT8** | TGCTGTTGGGATAGCGAAGG | CACAAATCCACACATATCATTAGGG |

**Table S5. Composition of internal standard mixture for lipidomics analysis.** Stock solutions of individual internal standards were prepared in the mixture of CHCl3/MeOH (1:1, v/v).

| Lipid class | Internal standard | Molecular Weight | Stock solution [mg/mL] | V [µL] | Concentration [pmol/mg of cells] |
|---|---|---|---|---|---|
| **SM** | **SM 18:1;O2/18:1-D9** | 737.6391 | 1 | 96 | 17.35 |
| | **SM 18:1;O2/12:0** | 646.5049 | 2 | 42 | 17.32 |
| **Cer** | **Cer 18:1-D7;O2/18:0** | 572.5868 | 1 | 12 | 2.79 |
| | **Cer 18:1;O2/17:0** | 551.5277 | 2 | 6 | 2.90 |
| | **Cer 18:1;O2/12:0** | 481.7900 | 2 | 5.1 | 2.82 |
| **GlcCer** | **GlcCer 18:1-D5;O2/18:0** | 732.6270 | 1 | 18 | 3.28 |
| | **GlcCer 18:1;O2/12:0** | 643.9350 | 0.2 | 79.5 | 3.29 |
| **GalCer** | **GalCer 18:1-D7;O2/13:0** | 664.5600 | 0.004 | 420 | 0.34 |
| | **GalCer 18:1;O2/12:0** | 643.5020 | 0.004 | 405 | 0.34 |
| **LacCer** | **LacCer 18:1-D7;O2/15:0** | 854.6455 | 1 | 6 | 0.94 |
| | **LacCer 18:1;O2/12:0** | 805.5551 | 2 | 3 | 0.99 |
| **Ga2Cer** | **Ga2Cer 18:1;O2/17:0** | 875.6300 | 0.025 | 3 | 0.01 |
| **Sulfatides** | **SHexCer 18:1-D7;O2/13:0** | 761.5500 | 0.005 | 1.2 | 0.001 |
| | **SHexCer 18:1;O2/12:0** | 740.4860 | 0.005 | 1.35 | 0.001 |
| **SPB** | **SPB 18:1-D7;O2** | 306.3264 | 0.2 | 9 | 0.78 |
| | **SPB 17:1;02** | 285.2668 | 0.4 | 4.5 | 0.84 |

**Table S6. Constructs used in this study.**

| Construct name | Amino acids |
|---|---|
| NF155 IgG1-Fn4 (FL ECD) | I25-G1043 |
| NF155 IgG1-4 | I25-S435 |
| NF155 Fn1-4 | R626-G1043 |
| NF155 IgG5-Fn2 | E528-D819 |
| NF155 IgG5-Fn4 | E528-G1043 |
| NF155 IgG1-Fn2 | I25-D819 |
| NF186 IgG1-Fn4 (FL ECD) | I25-G1109 |
